# Zinc is a Key Regulator of the Sperm-Specific K^+^ Channel (Slo3) Function

**DOI:** 10.1101/2024.12.12.628223

**Authors:** Rizki Tsari Andriani, Tanadet Pipatpolkai, Haruhiko Miyata, Masahito Ikawa, Yasushi Okamura, Takafumi Kawai

## Abstract

The voltage- and pH-gated Slo3 potassium channel is exclusively expressed in mammalian spermatozoa. Its sensitivity to both voltage and alkalization plays a crucial role in sperm fertility, which is tightly coupled to the capacitation process. Here we show that sperm-enriched divalent cation Zn^2+^ undergoes dynamic alteration in spermatozoa during capacitation. We also found that intracellular Zn^2+^ regulates alkalinization-induced hyperpolarization in mouse spermatozoa which is mediated by Slo3 channel. Further examination of zinc regulation in mouse Slo3 (mSlo3) revealed that, in *Xenopus* oocyte expression system, intracellular zinc directly inhibits mouse Slo3 currents in dose-dependent manner at micromolar concentrations, with exceptionally slow dissociation. By combining MD simulations and electrophysiology, we also identified amino acid residues contributing to the Zn^2+^ slow dissociation from Slo3 channels. Our studies uncover the importance of intracellular zinc dynamics and its regulatory role in ion channels during sperm capacitation.

## Introduction

Zinc is an essential metal ion involved in numerous biological functions, including as a regulatory component to various proteins such as enzymes, transcription factors, and ion channels (Fayyazuddin et al., 2000; Gao et al., 2017; Maret, 2013; Sun et al., 2007; Vallee & Falchuk, 1993; Zheng et al., 2001). In the reproductive system, spermatozoa is a type of cell that is known to have high intracellular zinc content, which influences its production, maturation, and physiological functions (Chia et al., 2000; Fallah et al., 2018; Henkel et al., 1999; Riffo et al., 1992; Yamaguchi et al., 2009). Zinc is also present in the seminal fluid at high concentrations (∼1-1.5 mM) (Henkel et al., 1999; Riffo et al., 1992), and extensive studies have elucidated their role in regulating sperm physiology. For instance, extracellular zinc has been shown to influence human sperm motility, which negatively correlates with the concentration (Henkel et al., 1999, 2005). Another study also suggested that increased zinc concentration inhibits sperm motility (Riffo et al., 1992). Despite considerable evidence that sperm motility is regulated by extracellular zinc, intracellular zinc has also been implicated in playing a regulatory role. One study demonstrated that chelating intracellular zinc reduces sperm motility and velocity (Sørensen et al., 1999). However, the underlying mechanism by which zinc regulates sperm motility remains unclear.

During fertilization, spermatozoa undergo capacitation, a series of physiological processes that determine the success of fertilizing the egg within the female reproductive tract (Austin, 1951; Chang, 1951). This capacitation involves the changes of activities in various ion channels, which leads to diverse physiological changes, such as pH, calcium level, and membrane potential (Arnoult et al., 1999; Breitbart, 2002; Publicover et al., 2007; Vredenburgh-Wilberg & Parrish, 1995; Y. Zeng et al., 1995, 1996). Moreover, recent studies have shown that the activity of ion channels is regulated by zinc. Indeed, extracellular zinc has been shown to inhibit the flagellar voltage-gated proton channel Hv1, which is supposed to control intracellular pH in human sperm (Grahn et al., 2023; Lishko et al., 2010; Zhao et al., 2018). Thus, it is possible that the high levels of zinc in spermatozoa regulates their physiological properties.

In the present study, we find that intracellular zinc is exported during capacitation, indicating that zinc dynamics in spermatozoa play an important role in fertilization. Furthermore, zinc significantly inhibited the alkalinization-induced hyperpolarization in mouse sperm, which is mediated by Slo3 (K_Ca_5.1), a sperm specific pH- and voltage-gated potassium channel (Schreiber et al., 1998; X. Zhang et al., 2006).

We further discovered that intracellular zinc directly inhibits mouse Slo3 (mSlo3) currents using *Xenopus* oocytes. Of note, this inhibition is pH-dependent, voltage-dependent and has a long-lasting inhibitory effect even after the washout. Coexpression of mSlo3 with the auxiliary subunit Lrrc52, to mimic the native environment (Yang et al., 2011; X. H. Zeng et al., 2015), consistently demonstrates zinc’s inhibitory effect. Additionally, we identify potential key amino acids in the mSlo3 channel that serve as the zinc binding site. Our findings highlight the importance of intracellular zinc dynamics and its regulatory role over ion channels during capacitation in spermatozoa.

## Results

### Intracellular zinc is exported from sperm during capacitation

To investigate the role of intracellular zinc during capacitation, we monitored the intracellular zinc concentration utilizing the zinc-sensitive fluorophore FluoZin3-AM. Fluorescence signals were successfully detected in both the head and flagellum of isolated mouse spermatozoa (Fig. 1A). As expected, the head of the spermatozoa showed a stronger fluorescence signal compared to the flagellum, likely due to its higher zinc capacity. We also treated spermatozoa with Toyoda, Yokoyama, Hoshi (TYH) medium (Toyoda et al., 1971), which is well established capacitation-inducing medium *in vitro*. We observed that there was a gradual and significant decrease in fluorescence intensity in both regions (Fig. 1B), particularly prominent in the flagellum (Fig. 1C). This decline suggests the active release of intracellular zinc from sperm flagellum occurs during capacitation. In contrast, the fluorescence intensity of the control group of non-capacitated sperm remained unchanged (Fig. 1B).

**Figure 1.**
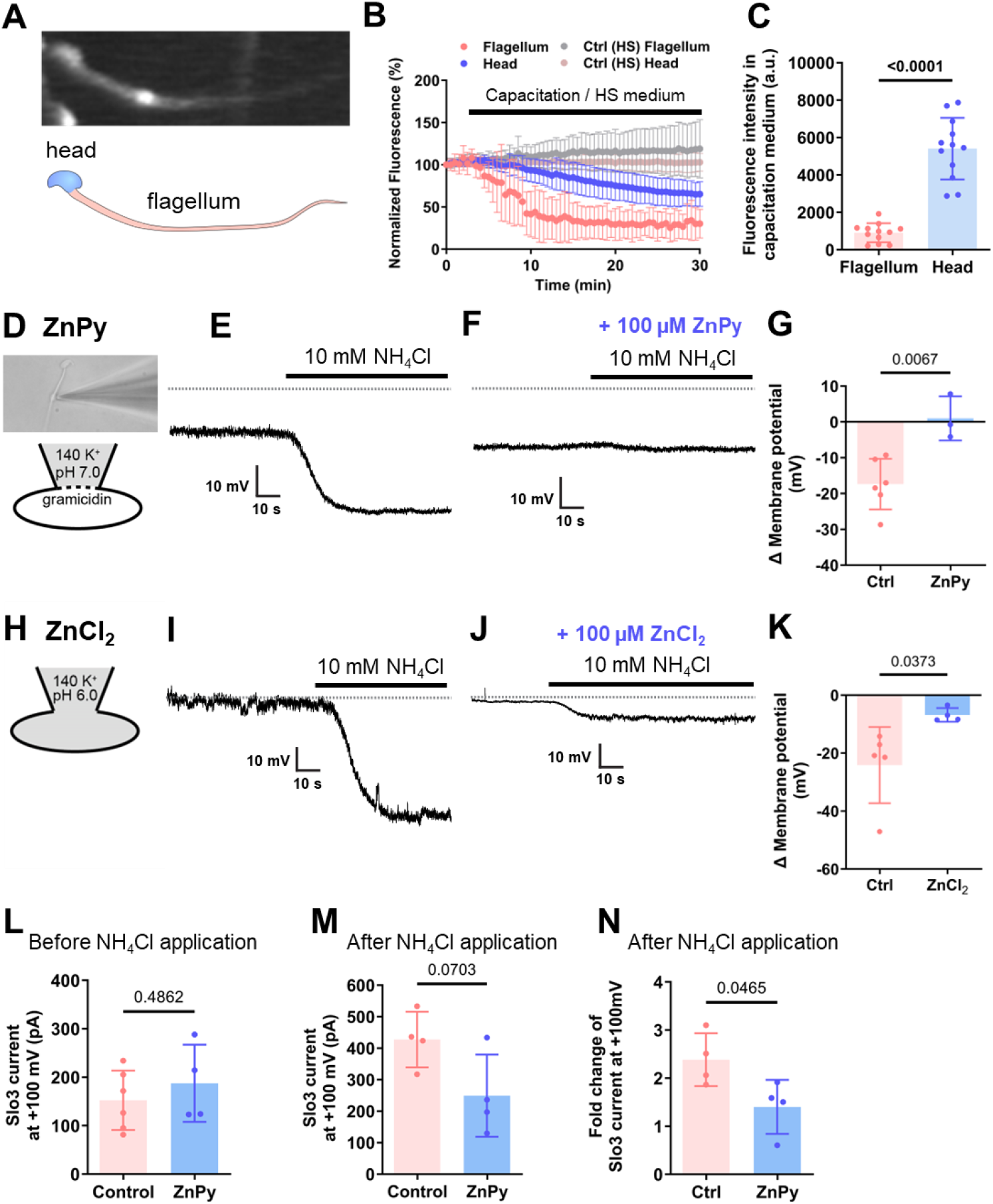
Intracellular zinc is exported from sperm during capacitation and inhibits the alkalinization-induced hyperpolarization in sperm. (**A-C**) Intracellular zinc is exported from sperm during capacitation. (**A**) Representative image of FluoZin3-AM fluorescence observed in spermatozoa (head and flagellum, as indicated). (**B**) Time course of normalized FluoZin3-AM fluorescence response upon the application of capacitation medium. Both head and flagellum showed a decreased fluorescence response, indicating that intracellular zinc was exported from sperm in response to the application of capacitation medium. (**C**) Comparison of the fluorescence intensity from flagellum (908.6 ± 508.2; n=12) and head (5409 ± 1648; n=12) after capacitation. Unpaired t-test, p<0.0001. All error bars are ± SD centered on the mean. (**D-H**) Intracellular zinc inhibits the alkalinization-induced hyperpolarization in sperm. (**D-G**) Membrane potential (Vm) recording by perforated patch-clamp were performed in mouse spermatozoa as depicted in (**D**). (**E**) Representative trace of Vm recording for control group. The application of NH_4_Cl increased the intracellular pH (pHi) which then hyperpolarized the membrane, the process called alkalinization-induced hyperpolarization. Application of NH_4_Cl alone hyperpolarized the membrane by -17.37 ± 7.09 mV (n=6). (**F**) Representative trace of Vm recording in the presence of zinc ionophore (zinc pyrithione, ZnPy). When 100 µM ZnPy was applied on the top of NH_4_Cl application, no hyperpolarization was observed (ΔVm = 0.99 ± 6.17 mV; n=3). All the dotted lines indicated 0 mV level. (**G**) Comparison of ΔVm recording between control (n=6) and in the presence of 100 µM ZnPy (n=3), p=0.0067, unpaired t-test. (**H-K**) Vm recording by whole-cell patch-clamp configuration in mouse spermatozoa as depicted in (**H**). (**I**) Representative trace of Vm recording for control group. The application of NH_4_Cl increased the [pHi] which then hyperpolarized the membrane (ΔVm= -24.12 ± 13.16 mV; n=5). (**J**) Representative trace of Vm recording in the presence of 100 µM ZnCl_2_. When zinc was applied to the intracellular part of spermatozoa on top of NH_4_Cl application, alkalinization-induced hyperpolarization was inhibited (ΔVm= -6.80 ± 2.35 mV; n=4). All the dotted line indicated 0 mV level. (**K**) Comparison of ΔVm recording between control (n=5) and in the presence of 100 µM ZnCl_2_ (n=4), p=0.0373, unpaired t-test. (**L-N**) Slo3 current recording using perforated patch-clamp in sperm. (**L**) Comparison of Slo3 current amplitude at +100 mV before NH_4_Cl application between control (152.4 ± 25.05 pA; n=6) and in the presence of 100 µM ZnPy (187.5 ± 79.53 pA; n=4), p=0.4862, Welch’s t-test. (**M**) Comparison of Slo3 current amplitude at +100 mV after NH_4_Cl application between control (427.6 ± 88.28 pA; n=4) and in the presence of 100 µM ZnPy (249.2 ± 130.6 pA; n=4), p=0.0703, Welch’s t-test. (**N**) Comparison of the fold change in Slo3 current amplitude at +100 mV after NH_4_Cl application between control (2.39 ± 0.55; n=4) and in the presence of 100 µM ZnPy (1.4 ± 0.56; n= 4), p=0.0465, Welch’s t-test. All error bars are ± SD centered on the mean.

Additionally, we examined the effect of intracellular zinc on sperm motility before and after capacitation (Fig. 1—figure supplement 1A-E). We incubated the isolated spermatozoa with cell permeable Zn^2+^ chelator N,N,N’,N’-Tetrakis(2-pyridylmethyl)ethylenediamine (TPEN) and measured the motility parameters before and after capacitation. We found that VAP (average path velocity), VCL (curvilinear velocity), and VSL (straight-line velocity) were influenced by the TPEN treatment only after the capacitation, as shown in Fig. 1—figure supplement 1. These results demonstrate that the dynamics of zinc levels during capacitation potentially contributes to sperm motility, highlighting the importance of zinc action in sperm physiology.

### Intracellular zinc inhibits the alkalinization-induced hyperpolarization in sperm

It is already known that sperm capacitation is well associated with the increase of intracellular pH (Vredenburgh-Wilberg & Parrish, 1995; Y. Zeng et al., 1996), which leads to the hyperpolarization of the membrane (Arnoult et al., 1999; Y. Zeng et al., 1995) as well as the elevation of intracellular Ca^2+^ concentration level (Breitbart, 2002; Publicover et al., 2007) through diverse ion channel activities. To explore whether these pathways are influenced by intracellular zinc, we used perforated patch-clamp techniques. It has been reported that under the whole-cell current clamp of mouse epididymal spermatozoa, resting membrane potential (Vm) is hyperpolarized after intracellular alkalinization (Navarro et al., 2007). Therefore, we applied 10 mM NH_4_Cl, which is routinely used to alkalize intracellular pH (Madshus, 1988), to spermatozoa. As expected, we observed that it hyperpolarized the membrane by -17.37 ± 7.09 mV; n=6 (Fig. 1E, G). Then, we applied 10 mM NH_4_Cl with 100 µM zinc pyrithione (ZnPy), a zinc ionophore. In contrast to the control group, there was no remarkable hyperpolarization in the presence of ZnPy as the ΔVm was 0.99 ± 6.17 mV; n=3 (Fig. 1F, G). These findings indicate that either extracellular or intracellular zinc inhibits the alkalinization-induced hyperpolarization in mouse sperm.

To examine the possibility that intracellular, but not extracellular, zinc is responsible for inhibiting the alkalinization-induced hyperpolarization, we measured membrane potential of sperm using the patch-clamp technique in conventional whole-cell configuration with pH_i_ = 6.0 (Fig. 1H). In the test group, the patch pipette contained 100 µM ZnCl_2_ (Fig. 1J), while the control group did not (Fig. 1I). Consistent with the perforated patch-clamp results, we observed alkalinization-induced hyperpolarization upon applying NH_4_Cl in the control (ΔVm= -24.12 ± 13.16 mV; n=5; Fig. 1I, K). In contrast, alkalinization-induced hyperpolarization was reduced in the 100 µM ZnCl_2_ group (ΔVm= -6.80 ± 2.35 mV; n=4; Fig. 1J, K).

We also measured Slo3 current using perforated patch-clamp recordings in spermatozoa treated with ZnPy, before and after the addition of NH_4_Cl. Control experiments were conducted in the absence of ZnPy, in which Slo3 current were recorded before and after the application of 10 mM NH_4_Cl (Fig. 1L-N; Fig. 1—figure supplement 2A, B). Slo3 current in sperm tended to be inhibited by zinc, as shown by the plot of absolute Slo3 current after the addition of 10 mM NH_4_Cl in the absence of ZnPy (control) and in the presence of 100 µM ZnPy (Fig. 1L, M). There was a decrease in the fold change calculated from the absolute current before and after the addition of 10 mM NH_4_Cl of ZnPy treated group (1.4 ± 0.56; n=4) compared to the control group (2.39 ± 0.55; n=4; Fig. 1N). Taken together, these results confirmed that intracellular zinc indeed inhibits alkalinization-induced hyperpolarization in mouse sperm.

### Zinc inhibits mouse Slo3 (mSlo3) current

Our findings suggest that intracellular zinc inhibits a key process in sperm capacitation, specifically the alkalinization-induced hyperpolarization. Previous studies have identified the pH-and voltage-dependent potassium channel Slo3 is responsible for the principal K^+^ current (I_KSper_) in mouse spermatozoa (Navarro et al., 2007; Santi et al., 2010; Schreiber et al., 1998; X. H. Zeng et al., 2011). During capacitation, the rise in pHi leads to the activation of Slo3 channels, resulting in membrane hyperpolarization (Santi et al., 2010). Given this context, we next investigated whether intracellular zinc acts directly on the Slo3 channel.

We expressed the mSlo3 channel in the *Xenopus* oocyte expression system and conducted inside-out patch-clamp recordings. The bath solution was initially set to pH 8.0, and recordings were performed using a high-speed perfusion system. A ramp pulse from -100 mV to + 100 mV with a holding potential of -60 mV was applied. We observed a robust pH- and voltage-dependent mSlo3 current at pH 8.0 (Fig. 2A, B), consistent with previous reports (Leonetti et al., 2012; Schreiber et al., 1998; X. Zhang et al., 2006). The application of 100 µM of ZnCl_2_ resulted in the inhibition of 36.62 ± 11.65 % (n=6) of mSlo3 current (Fig. 2A, B). Of note, the inhibited mSlo3 current by zinc was never restored after the wash out of zinc even for more than 30 seconds. However, when pH 6.0 was applied to completely abolish the mSlo3 current, followed by the application of pH 8.0, the mSlo3 current recovered (Fig. 2A, B). We also determined the IC_50_ of zinc inhibition by applying zinc at concentrations 0.1, 1, 10, 100, and 1000 µM for 30 seconds. The dose-response curve, fitted to the Hill equation with a standard slope (Hill slope of -1.0), yielded an IC_50_ = 145.2 µM (Fig. 2C).

**Figure 2.**
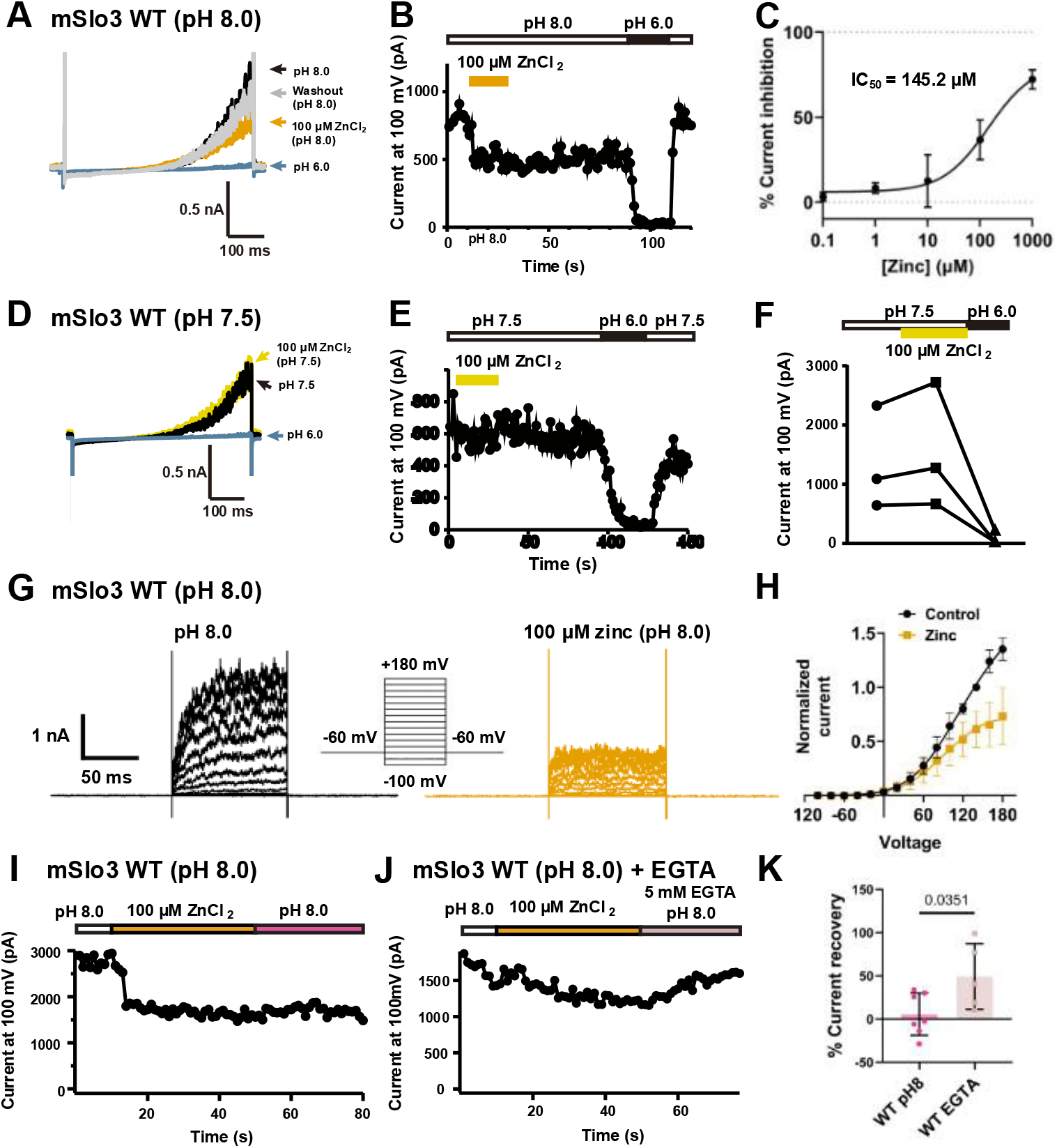
Zinc inhibits mouse Slo3 (mSlo3) current. (**A-C**) Inside-out patch-clamp recording of mSlo3 with pHi=8.0 by applying ramp pulse from -100 mV to +100 mV with the holding potential of 0 mV. (**A**) Representative current traces upon the application of pH 8.0 (black); 100 µM of zinc in pH 8.0 (yellow orange); pH 6.0 (blue); and wash-out by pH 8.0 (gray) after pH 6.0. (**B**) Time course of the change in current amplitude at +100 mV in response to pH 8.0, 100 µM zinc, pH 6.0, and pH 8.0 as indicated (white: pH 8.0; yellow orange: 100 µM zinc in pH 8.0; solid black: pH 6.0). (**C**) Dose-response curve of zinc inhibition in mSlo3 current (pH 8.0). % current inhibition was plotted assuming that the dose response curve has a standard slope, equal to a Hill slope of - 1.0. IC_50_ = 145.2 µM; n = 2-6. (**D-F**) Inside-out patch-clamp recording of mSlo3 in *Xenopus* oocyte with pHi=7.5 by applying ramp pulse from –100 mV to +100 mV with the holding potential of 0 mV. (**D**) Representative current traces upon the application of pH 7.5 (black); µM of zinc (yellow); and pH 6.0 (blue). (**E**) Time course of the change in current amplitude at +100 mV in response to pH 7.5, 100 µM zinc, and pH 6.0 as indicated (white: pH 7.5; yellow: 100 µM zinc in pH 7.5; solid black: pH 6.0). (**F**) Current amplitude at +100 mV upon the application of pH 7.5, 100 µM zinc in pH 7.5, and pH 6.0 from individual cell, as indicated. (**G-H**) Inside-out patch-clamp recording of mSlo3 with pHi=8.0 by applying step pulses from –100 mV to +180 mV with the holding potential of -60 mV. Zinc inhibition is voltage-dependent. (**G**) Representative current traces upon the application of pH 8.0 (black) and 100 µM of zinc in pH 8.0 (yellow orange). (**H**) I-V relationship of control (pH 8.0; black) and zinc (100 µM; yellow orange). Normalization was performed by dividing the absolute current of mSlo3 at pH 8.0 of each voltage by the absolute current at the pre-determined highest voltage that still produced a stable mSlo3 current (i.e., good patch, good clamp). In this analysis, +140 mV was chosen as the highest voltage for normalization, since in few cells the patch was lost at +160mV and +180mV. Similar to the control condition, the absolute current of mSlo3 in the presence of 100 µM zinc was normalized to the absolute current of the control at +140 mV. I-V curve was fitted by Boltzmann equation for control and followed by Woodhull equation for zinc inhibition (V_50 Ctrl_ = 85.24; Slope _Ctrl_ = 35.51; V_block_ Zn = 188.8; Slope_block_ Zn = - 115.3; n=13). (**I-K**) Inside-out patch-clamp recording of mSlo3 with the application of 5 mM EGTA in pH=8.0 by applying ramp pulse from –100 mV to +100 mV with the holding potential of 0 mV. Zinc has a long-lasting inhibitory effect on mSlo3, and it could be rescued by EGTA (**I**) Time course of the change in current amplitude at +100 mV in response to pH 8.0, 100 µM zinc, and pH 8.0 as indicated (white: pH 8.0; yellow orange: 100 µM zinc in pH 8.0; pink: pH 8.0 for wash-out). (**J**) Time course of the change in current amplitude at +100 mV in response to pH 8.0, 100 µM zinc, and 5 mM EGTA in pH 8.0 as indicated (white: pH 8.0; yellow orange: 100 µM zinc in pH 8.0; light lilac: 5 mM EGTA in pH 8.0 for wash-out). (**K**) Comparison of percentage of current recovery between pH 8.0 only (5.62 ± 24.40 %; n= 7) and 5 mM EGTA in pH 8.0 (49.22 ± 37.98 %; n=5), p=0.0351, unpaired t-test. All error bars are ± SD centered on the mean.

We also tested the effect of zinc at a lower pH of 7.5. We observed the pH- and voltage-dependent mSlo3 current at this pH. However, the 100 µM of ZnCl_2_ application did not affect the mSlo3 current (Fig. 2D-F). To rule out instrumental and human error, we also tested the effect of pH 6.0 which suppresses Slo3 activity. As expected, the application of pH 6.0 completely inhibited the mSlo3 current, and the subsequent application of pH 7.5 restored the current (Fig. 2D-F).

We also conducted step pulse protocols from -100 mV to +180 mV with a holding potential of -60 mV to assess whether zinc inhibition is voltage-dependent. At pH 8.0, we observed a robust mSlo3 current, which was inhibited by 100 µM ZnCl_2_ (Fig. 2G, H). The current-voltage (I-V) relationship indicated voltage-dependent zinc inhibition (Fig. 2H). The ratio of I_zinc_ to I_control_ plotted against membrane potential also demonstrated that zinc inhibition of mSlo3 current is voltage-dependent (Fig. 2—figure supplement 1B).

In addition, we plotted the conductance-voltage (G-V) relationship by calculating the conductance from the maximum current amplitude at steady-state, as the tail current recordings of mSlo3 were unsuccessful. We noted that V_50_ of mSlo3 current shifted to more hyperpolarized potentials in the presence of zinc (V_50_ control = 80.67 ± 2.32; V_50_ zinc = 49.68 ± 1.38; n=13; Fig. 2—figure supplement 1C).

As we observed the long-lasting inhibitory effect of zinc on mSlo3 current, we examined whether a zinc chelator could restore the mSlo3 current to the baseline level without applying acidic pH. We used 5 mM EGTA, a divalent cation chelator which strongly chelates zinc (Kd_(Zn)_ ∼1 nM) (Hu et al., 2009). Zinc-inhibited mSlo3 current could not be recovered by pH 8.0 alone (Fig. 2I). However, significant current recovery was observed when 5 mM EGTA was included in the pH 8.0 bath solution (Fig. 2J). Specifically, 5 mM EGTA recovered 49.22 ± 37.98 % of the mSlo3 current (n=5), compared to pH 8.0 alone, which showed no significant recovery (5.62 ± 24.4 %; n= 7), as shown in Fig. 2K.

Additionally, we investigated the presence of endogenous zinc in Slo3 by perfusing pH 8.0 followed by 5 mM EGTA at the beginning of the recording to detect any changes in the baseline current. The baseline current showed a slight increase upon perfusion with 5 mM EGTA, indicating the presence of endogenous zinc affecting the baseline current in Slo3 (Fig. 2—figure supplement 1D, E).

Taken together, these results showed that zinc has a long-lasting inhibitory effect on mSlo3 current that cannot be reversed by pH 8.0 alone. The inhibitory effect is pH-dependent, as complete inhibition at pH 6.0 followed by recovery at pH 8.0 was observed. In addition, zinc inhibition is voltage-dependent, and the application of 5 mM EGTA, which strongly chelates zinc, can partially recover mSlo3 current.

### Zinc more efficiently inhibits mSlo3 current when coexpressed with the auxiliary subunit Lrrc52

In its native environment of spermatozoa, Slo3 is expressed with its auxiliary subunit Lrrc52 (Yang et al., 2011; X. H. Zeng et al., 2015). Lrrc52 shifts the Slo3 gating to more negative potentials, making the channel easier to open (Yang et al., 2011). We coexpressed mSlo3 with mouse Lrrc52 (mLrrc52) in the *Xenopus* expression system to perform inside-out patch clamp recordings.

The mSlo3+mLrrc52 current was observed upon the application of pH 8.0. A ramp pulse protocol from -100 mV to +100 mV was applied with a holding potential of 0 mV. Similar to mSlo3 alone, in the presence of mLrrc52, mSlo3 current was also inhibited by 100 µM ZnCl_2_ (Fig. 3A, B). In addition, zinc inhibition was not recovered after washing with the zinc-free solution (Fig. 3B). A significant leftward shift in IC_50_ was observed when the dose-response curve was plotted for coexpression with mLrrc52 (Fig. 3C; IC_50_ = 15.2 µM; n= 3-8) compared to mSlo3 alone (Fig. 2C, Fig. 3C).

**Figure 3.**
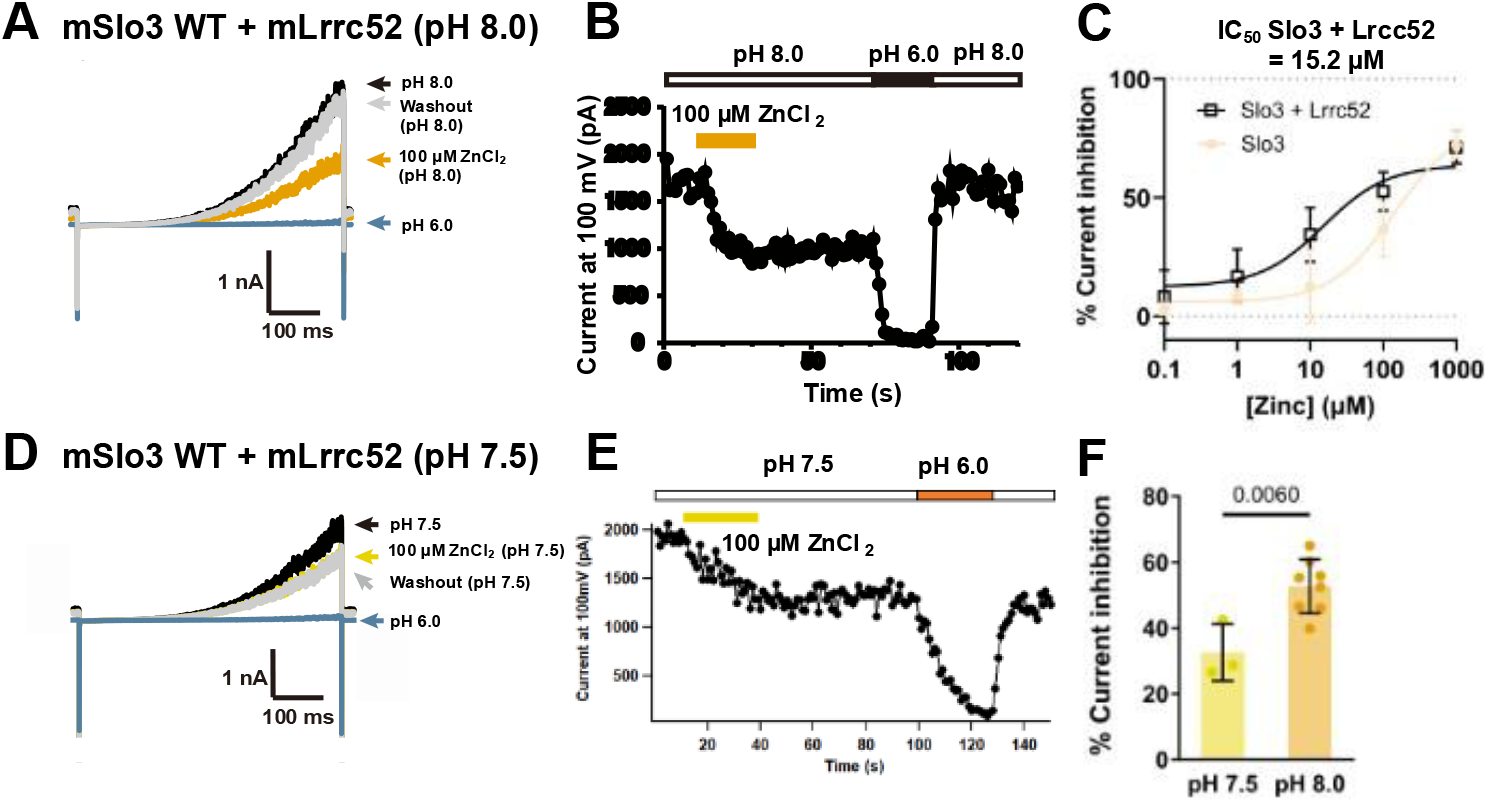
Zinc inhibits mSlo3 current when co-expressed with gamma subunit Lrrc52. (**A-B**) Inside-out patch-clamp recording of mSlo3 co-expressed with mLrrc52 with pHi=8.0 by applying ramp pulse from –100 mV to +100 mV with the holding potential of 0 mV. (**A**) Representative current traces upon the application of pH 8.0 (black); 100 µM of zinc in pH 8.0 (yellow orange); pH 6.0 (blue); and wash-out by pH 8.0 after pH 6.0 (gray). (**B**) Time course of the change in current amplitude at +100 mV in response to pH 8.0, 100 µM zinc, pH 6.0, and pH 8.0 as indicated (white: pH 8.0; yellow orange: 100 µM zinc in pH 8.0; solid black: pH 6.0). (**C**) Dose-response curve of zinc inhibition in mSlo3 co-expressed with mLrrc52 current (pH 8.0). % current inhibition was plotted assuming that the dose response curve has a standard slope, equal to a Hill slope of -1.0. IC_50_ = 15.2 µM; n = 3-8. (**D-E**) Inside-out patch-clamp recording of mSlo3 in *Xenopus* oocyte co-expressed with mouse Lrrc52 (mLrrc52) with pHi=7.5 by applying ramp pulse from –100 mV to +100 mV with the holding potential of 0 mV. (**D**) Representative current traces upon the application of pH 7.5 (black); µM of zinc (yellow); pH 7.5 for wash-out (grey); and pH 6.0 (blue). (**E**) Time course of the change in current amplitude at +100 mV in response to pH 7.5, 100 µM zinc, pH 6.0, and pH 7.5 for wash-out as indicated (white: pH 7.5; yellow: 100 µM zinc in pH 7.5; orange pH 6.0). (**F**) Comparison of the percentage of current inhibition by 100 µM zinc between pH 7.5 (32.51 ± 8.62 %; n=3) and pH 8.0 (52.61 ± 8.23 %; n=8), p=0.006; unpaired t-test. All error bars are ± SD centered on the mean.

Additionally, when pH 7.5 was applied to depolarized potentials, mSlo3+mLrrc52 current was activated and in contrast with mSlo3 alone (Fig. 2D-F), 100 µM ZnCl_2_ already inhibited mSlo3+mLrrc52 current at pH 7.5 (Fig. 3D, E, Fig. 3—figure supplement 1D). Alkalinization appeared to increase the percentage of current inhibition by 100 µM ZnCl_2_ (Fig. 3F; pH 7.5 = 32.51 ± 8.62 %; n=3 and pH 8.0 = 52.61 ± 8.23 %; n=8).

We next conducted recordings using a step pulse protocol from -100 mV to +180 mV with a holding potential of -60 mV. Upon applying pH 8.0, we observed a significant mSlo3+mLrrc52 current, which was inhibited by 100 µM ZnCl_2_ (Fig. 3—figure supplement 1A, B). Again, the I-V relationship confirmed that zinc inhibition was voltage-dependent (Fig. 3—figure supplement 1B). V_50_ of mSlo3+mLrrc52 current was also shifted into more hyperpolarized potentials in the presence of zinc (V_50_ control = 72.93 ± 2.06 V_50_ zinc = 56.27 ± 2.75; n=13; Fig. 3—figure supplement 1C).

Taken together, when co-expressed with mLrrc52, mSlo3 current was more efficiently inhibited by zinc.

### E169 and E205 located within VSD are important for the sustained zinc inhibition of mSlo3 current

To elucidate the mechanism of zinc inhibition on the mSlo3 channel, we aimed to identify the structural determinants required for this inhibition using a range of computational tools and mutagenesis. First, we used the AlphaFold3 structure prediction webserver (Abramson et al., 2024) to predict the zinc binding site on the mSlo3 channel. Notably, the prediction has shown that the majority of the zinc binding sites are located in the channel’s intracellular regulator of K^+^ conductance (RCK) domain, which consists of RCK1 and RCK2, with very few zinc sites in the transmembrane domain (Fig. 4—figure supplement 1A). The RCK domain is also recognized as crucial for zinc binding in Slo channel families (Hou et al., 2010; J. Zhang et al., 2023).

**Figure 4.**
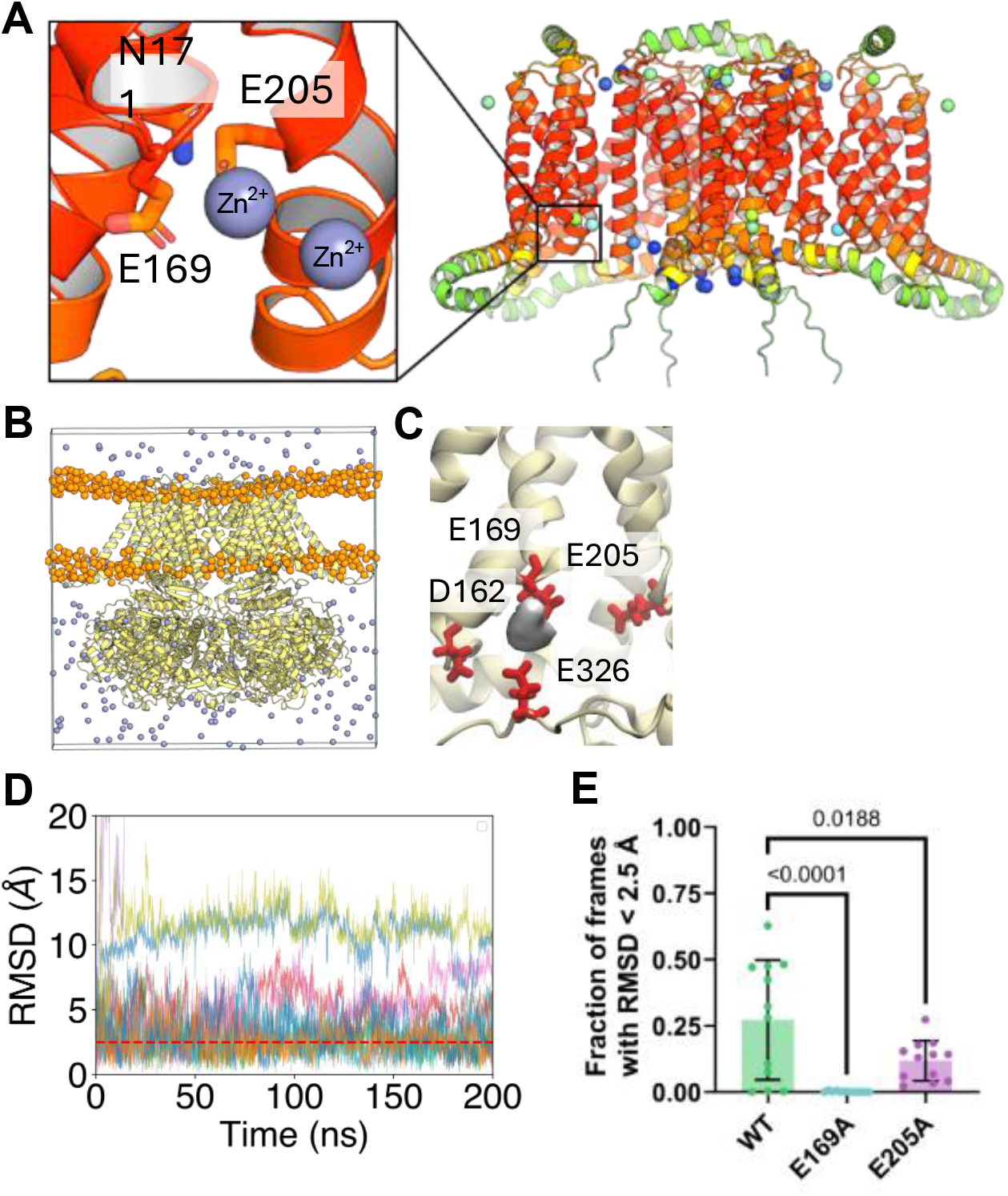
Computational studies show that the Zn binding site on mSlo3 is located near E169 and E205. (**A**) AlphaFold3 prediction of the Zn binding site on the transmembrane segment of the Slo3 channel. Zn is shown as grey spheres, and Slo3 is shown using an orange cartoon. (**B**) The system is set up for flooding simulations. The phosphorus atoms on the POPC lipid are shown in orange. All Zn ions in the system are shown as grey spheres. The Slo3 channel is shown in the yellow cartoon. All water, KCl and other atoms within POPC molecules are omitted for clarity. (**C**) The density of Zn ions within 4 Å of the protein molecules was calculated using VolMap averaged across 200 ns flooding simulations (n = 3). Key acidic contacting residues are shown as red sticks. (**D**) Root mean square (RMSD) calculation of the 12 Zn ions fitted to the C alpha atoms of the protein on the first frame for 200 ns. Given that there are 4 Zn in each subunit per repeat, this leads to 12 data sets shown in different colored plots. The red dotted line marked the cutoff at 2.5 Å. (**E**) The fractions of frames where the RMSD of zinc ions are less than 2.5 Å in WT channel, E169A and E205A mutants (p<0.0001, p=0.0188, respectively). Statistical analysis was done by one-way ANOVA with Dunnett’s post-hoc test compared to the WT group for each mutant (n=12). All error bars are ± SD centered on the mean.

To validate the site within the intracellular region, we conducted site-directed mutagenesis on intracellular domain focusing on RCK1, RCK1/RCK2 linker, and RCK2 (H354R, H383R, H423R, H516R, H564R, H606A, H695R, and H720R) to assess changes in channel sensitivity to zinc. However, the percentage of current inhibition varied across the mutated constructs, showing either increases or no appreciable change (Fig. 4—figure supplement 1B, C). We also generated a chimera where RCK2 of mSlo3 was replaced with RCK2 from mSlo1, but this chimera exhibited slightly increased sensitivity to zinc (Fig. 4—figure supplement 1B, C). Taken together, this concludes that, unlike Slo1, the zinc binding site is unlikely to be located in the RCK or the intracellular domain of the channel.

We then diverge our focus to the intracellular site of the channel. Given that zinc has been shown to shift the G-V relationship of the mSlo3 channel and its complex with Lrrc52 (Fig. 2—figure supplement 1C, Fig. 3—figure supplement 1C), we suspect that the binding site is more likely to be located in the transmembrane (TM) region. To enhance the prediction within the transmembrane segment, we computationally truncated the intracellular domain, allowing a key binding site to be found near E169 and E205 within the voltage-sensing domain (VSD) with the AlphaFold3 structure prediction webserver (Fig. 4A). Next, we conducted triplicates of 200 ns molecular dynamics simulations of the AlphaFold3 full-length Slo3 tetrameric structure to validate this binding site where the Slo3 channel is solvated in 0.15 M ZnCl_2_ and 0.15 M KCl (Fig. 4B). Here, we show that using MD simulations, a similar zinc site is also observed near E169 and E205, validating the prediction results by AlphaFold3 (Fig. 4C). In addition, we also observed another zinc binding near the pore at E310 and E313 (Fig. 5—figure supplement 1A).

**Figure 5.**
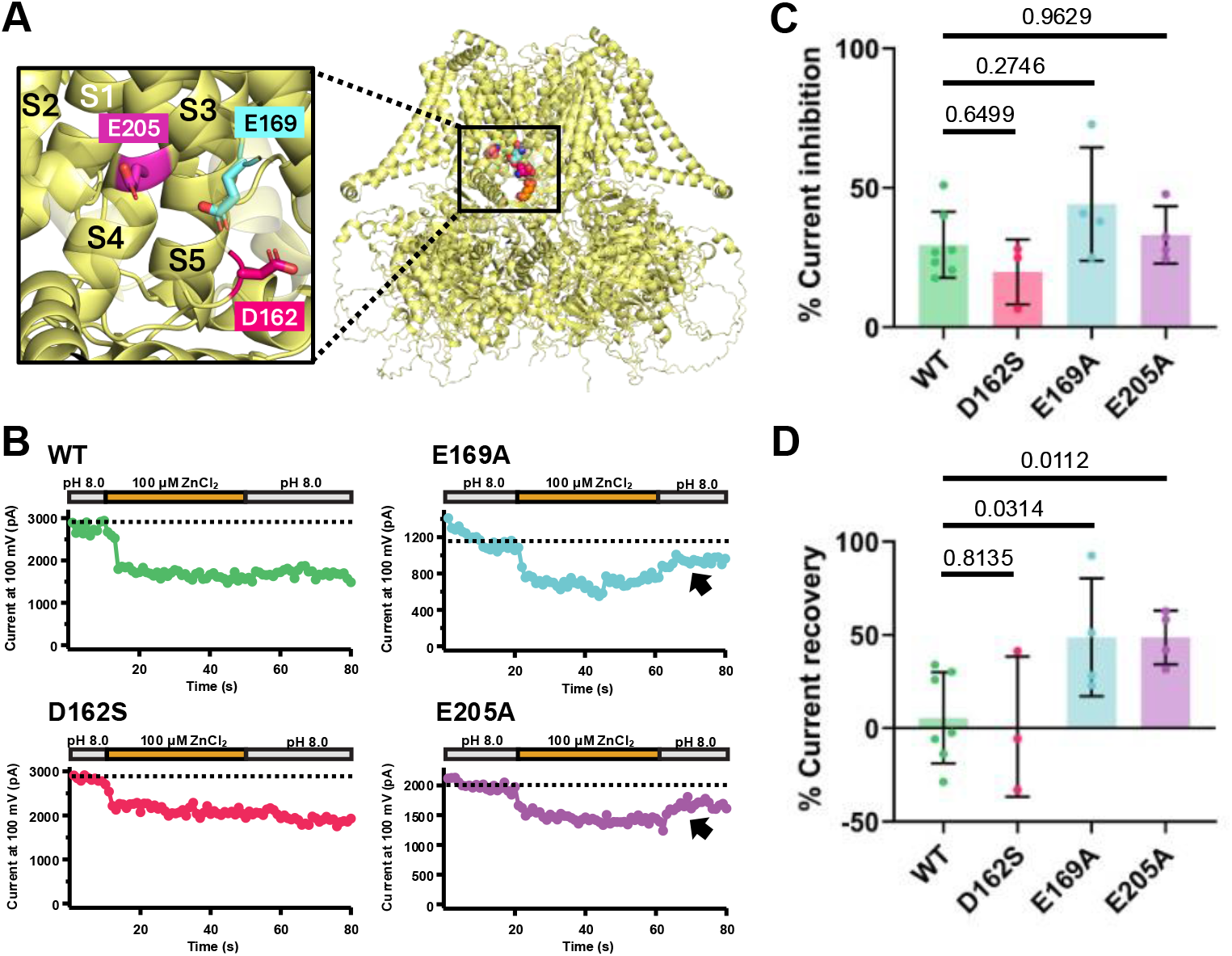
E169 and E205 located within VSD are important for the sustained zinc inhibition of mSlo3 current. (**A**) Tetrameric mSlo3 structure representation. Inset shows rotated view of selected region which indicates predicted zinc binding site from MD simulations located within the voltage-sensor domain: D162 (S2-S3 linker), E169 (S3 domain), and E205 (S4 domain) which have been mutated to elucidate the zinc binding site in mSlo3. mSlo3 structure representation in (**A**) was based on structure prediction using AlphaFold3 (Abramson et al., 2024), mSlo3 sequence was retrieved from NCBI NP_032458.3. Some portions of the S2 and S3 domains have been rendered highly transparent to enhance clarity. (**B**) Time course of the change in current amplitude at +100 mV in response to pH 8.0, 100 µM zinc, and pH 8.0 for wash-out as indicated for: WT, D162S, E169A, and E205A. The dotted line indicated the level of activated mSlo3 current at pH 8.0. Black arrows indicated the current recovery observed upon wash-out. (**C**) Comparison of the percent current inhibition upon the application of 100 µM zinc between WT, D162S, E169A, and E205A (p=0.6499, p=0.2746, p=0.9629, respectively). Statistical analysis was done by one-way ANOVA with Dunnett’s post-hoc test compared to the control group (WT), n=3-7 (**D**) Comparison of percentage of current recovery upon wash-out using pH 8.0 between WT, D162S, E169A, and E205A (p=0.8135, p=0.0314, p=0.0112, respectively). Statistical analysis was done by unpaired t-test compared to the WT group for each mutant, n=3-7. All error bars are ± SD centered on the mean.

To test the stability of zinc within E169 and E205, we removed all the ZnCl_2_ in the simulation box, leaving just 4 zinc ions near E169 residues and conducted three separate simulations for 200 ns, leading to 12 data points. Our simulations show that zinc remains stable, with the low root mean square deviation (RMSD) within its binding site in the transmembrane region, validating the stability within its binding site (Fig. 4D).

To validate the binding site further, we conducted a computational site-directed mutagenesis to further assess zinc stability after the mutation of the nearby acidic residue, E169A and E205A. Here, we show that the RMSD of the E169A mutant increases more than WT and E205A (Fig. 4E). Together, our simulation suggested that E169 and E205 are indeed important for the binding of zinc by Slo3.

We then examined zinc sensitivities of mSlo3 currents from oocytes expressing mutants by electrophysiology with mutation of zinc binding residues on the mSlo3 channel predicted by MD simulations, which is located in the S2-S3 linker (D162), S3 domain (E169), S4 domain (E205) and the intracellular linker between TM and RCK1 (E326) (Fig. 4C-E, Fig. 5A, Fig. 5—figure supplement 1B). All of these mutants exhibited similar current inhibition by zinc compared to wildtype (WT = 29.58 ± 11.84 %, n=7; D162S = 19.85 ± 11.63 %, n=3; E169A = 44.17 ± 20.27 %, n=4; E205 = 33.08 ± 10.28 %, n=4; E326 = 27.54 ± 1.27 %, n=2; Fig. 5B, C, Fig. 5—figure supplement 1C, D).

On the other hand, when we analyzed the percentage of current recovery of all the mutants, E169A and E205A showed significant current recovery upon the wash-out by pH 8.0 alone (WT = 5.62 ± 24.40 %, n=7; D162S = 0.86 ± 37.60 %, n = 3; E169A = 48.74 ± 31.61 %, n=4; E205A = 48.65 ± 14.37 %, n=4; E326 = -3.27 ± 1.43 %, n=2; Fig. 5B, D, Fig. 5— figure supplement 1C, E). Additionally, we performed sequence alignment by using ClustalO between mSlo3, mSlo1, and mSlo2.2. It is worth noting that only human and frog variants of Slo2.1 sequence are available in the database, so we included only Slo2.2 subtype, as our focus was on Slo3 in mouse sperm. Based on the alignment, E169 (mSlo3 numbering) is conserved among the Slo family channels in mice, while in contrast E205 (mSlo3 numbering) is not (Figure 5 – supplementary figure 1F).

Taken together, consistent with MD simulation results, our electrophysiological recordings demonstrated that the long-lasting inhibitory effect of zinc was partly abolished by these mutations. Thus, our findings highlight the contribution of E169A in the bottom of S3 domain and E205A in the bottom of S4 domain in zinc-mediated inhibition of mSlo3 current.

### Zinc alters the dynamics of VSD motion in mSlo3

Given the observed zinc-induced shift in the G-V relationship of both mSlo3 and mSlo3+mLrrc52, along with MD simulations and mutagenesis data, it is likely that VSD is involved in zinc inhibition of mSlo3. Then, we performed voltage-clamp fluorometry (VCF) experiments to track the motion of the VSD in the presence and absence of zinc. First, to identify suitable site for labeling with TMRM, we performed VCF scanning by introducing Cys mutations one at a time at the top of S3, S3-S4 linker, and the top of S4 of mSlo3 VCF construct (Fig. 6—figure supplement 1A). Two native, extracellular cysteines were mutated to avoid non-specific labeling (mSLo3 C19S/C131S) and the desired Cys mutations were then added on this double-mutant background. Initial scanning of VCF constructs L188C, K189C, S190C, and N191C (Fig. 6—figure supplement 1A-E) conjugated with TMRM resulted in no detectable fluorescence intensity (F) changes (ΔF). Subsequently, we coexpressed the mSlo3 VCF constructs with mLrrc52 to increase the level of mSlo3 expression in heterologous expression system (Leonetti et al., 2012). Out of three mutants coexpressed with mLrrc52 (L193C, G194C, and L195C; Fig. 6—figure supplement 1F-H), L193C, which is located in the top of S4 domain, produced fluorescence intensity changes in response to voltage steps (Fig. 6—figure supplement 1F) (ΔF/F= 1.15 ± 0.48 %; n=4).

**Figure 6.**
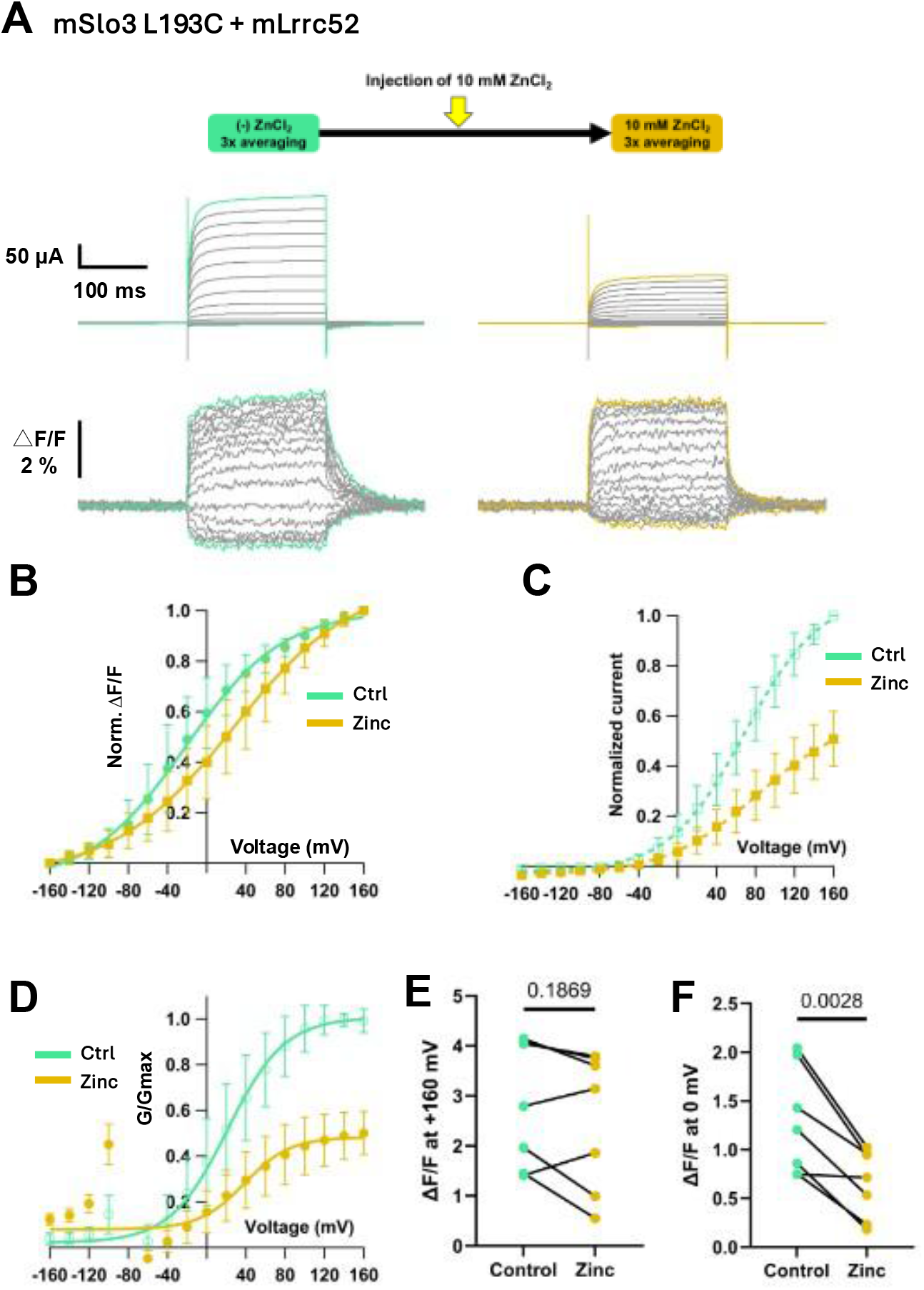
Zinc influenced the motion of the voltage-sensing domain (VSD) of mSlo3 channel. (**A**) Representative current traces and fluorescence signals of mSlo3 L193C coexpressed with Lrrc52 in the absence (green) and presence (yellow orange) of zinc. The inset illustrates a schematic timeline of direct zinc injection using a glass needle and positive pressure. Voltage steps from -160 mV to + 160 mV were applied with a holding potential of -60 mV. Zinc was injected into the same cell after the initial VCF recording (green), followed by a second VCF recording 60 seconds post-injection (yellow orange). ΔF/F_Control_ = 2.84 ± 1.25 % (n=7); ΔF/F_Zinc_ = 2.53 ± 1.37 % (n=7). The green and yellow orange traces represent the current and fluorescence signals at in +160 mV and -160 mV, in the absence (green) and presence (yellow orange) of zinc. (**B**) Comparison of the F-V relationship between control (green) and zinc (yellow orange). The F-V curve was fitted using Boltzmann equation (V_50_ control = =-21.68 ± 3.04; V_50_ zinc = =27.04 ± 1.05; n=7). Normalization was done based on the maximum ΔF/F for each control and zinc-injected conditions. (**C**) I-V relationship showing inhibition of mSlo3 L193C+Lrrc52 currents by direct zinc injection. Traces compare control (green) and zinc (yellow orange). (**D**) G-V relationship comparison of mSlo3 L193C current co-expressed with mLrrc52 between control (green circle) and zinc-injected (yellow orange circle) conditions (V_50_ control = 21.11 ± 4.8; V_50_ zinc = 38.62 ± 28.8; n=7). Normalization was done based on the maximum conductance in control. (**E**) Comparison of ΔF/F in the absence and presence of zinc at +160 mV from the same cell (p=0.1869, paired t-test, n=7). (**F**) Comparison of ΔF/F in the absence and presence of zinc at 0 mV from the same cell (p=0.0028, paired t-test, n=7). All error bars are ± SD centered on the mean.

We next performed VCF recordings in the absence and presence of zinc. Oocytes were held at -60 mV and the step pulses were applied from -160 mV to + 160 mV. Fluorescence change could be observed in response to voltage steps (ΔF/F_Control_ = 2.84 ± 1.25 % at +160 mV; n=7; Fig. 6A). A puff of 10 mM zinc solution was then manually injected into the same oocyte using a glass needle with positive pressure. 60 seconds post-injection, a second set of step pulses was applied to record the response under zinc application (ΔF/F_zinc_ = 2.53 ± 1.37 % at +160 mV; n=7; Fig. 6A). In the presence of zinc, the V_50_ of fluorescence-voltage (F-V) relationship was shifted into more depolarized potentials (V_50_ control = -21.68 ± 3.04; V_50_ zinc = 27.04 ± 1.05; Fig. 6B). This decrease in ΔF/F was more pronounced at certain voltages, such as 0 mV (Fig. 6E-F), and extended up to +80 mV. I-V relationship (Fig. 6C) showed that mSlo3 current was inhibited by zinc, consistent with what has been observed in the patch-clamp recordings (Fig. 2 and Fig. 3). Consistent with F-V relationship, the conductance-voltage (G-V) was also shifted into more depolarized potentials (V_50_ control = 21.11 ± 4.8; V_50_ zinc = 38.62 ± 28.8; Fig. 6D). Collectively, these results clearly indicate that the VSD motion of mSlo3 is influenced by the application of intracellular zinc.

## Discussion

The present study aims to understand the roles of intracellular zinc in sperm physiology, focusing on sperm capacitation. Our results demonstrated that intracellular zinc is exported from spermatozoa during capacitation. We further discovered that intracellular zinc regulates alkalinization-induced hyperpolarization in mouse spermatozoa, mediated by Slo3 channel. Detailed analysis of zinc’s regulatory effects mSlo3 in the *Xenopus* oocyte expression system revealed that intracellular zinc directly inhibits mSlo3 currents with exceptionally slow dissociation. By combining MD simulations and electrophysiology, we identified amino acid residues important for zinc-mediated inhibition of Slo3 activities.

### Zinc levels are dynamic during capacitation

Using FluoZin3-AM for zinc imaging, we confirmed the presence of intracellular zinc in sperm (Fig. 1A), which is consistent with previous findings (Henkel et al., 1999). Our observations revealed that treatment with capacitation medium induced a decrease in zinc fluorescence intensity (Fig. 1B, C), suggesting that zinc levels are dynamic during capacitation. Previous studies using boar, bull and human spermatozoa have identified four distinct patterns of zinc localization associated with the state of capacitation (Kerns et al., 2018). In non-capacitated sperm, high concentration of zinc is present throughout the whole sperm from the sperm head to the whole sperm-tail, and the subcellular zinc localization shifts to a medium level in both the sperm head and the sperm tail mid piece (Kerns et al., 2018). In congruent with this, our findings also demonstrated a decrease in fluorescence intensity in both the head and flagellum of mouse spermatozoa during capacitation (Fig. 1B, C). These results suggest that zinc dynamics appear to play an important role in sperm capacitation.

Indeed, we observed that zinc chelator significantly affected the sperm motility specifically in VAP (average path velocity), VCL (curvilinear velocity), and VSL (straight-line velocity) only after, but not before, capacitation (Fig. 1—figure supplement 1). Of note, it has been recently reported that all these motility parameters (VAP, VCL, and VSL) are reduced by Slo3-specific inhibitors in human sperm (M. Lyon et al., 2023). These findings are consistent with the idea that endogenous zinc dynamics control sperm motility through Slo3 during the capacitation process.

The identity of the transporters directly responsible for zinc dynamics remains unclear. To maintain zinc homeostasis, two protein families facilitate the transport of Zn^2+^ across cellular and intracellular membranes in opposite directions: zinc transporter proteins (ZnT) and Zrt-, Irt-like proteins (ZIP) (Kambe et al., 2004). ZIP12 has been reported to be highly expressed in mouse testis (Zhu et al., 2022), as well as ZnT-1 (Elgazar et al., 2005). The mechanisms of zinc transport in sperm during capacitation remains a subject to be investigated in future studies.

### Intracellular zinc directly inhibits mSlo3 channels

Our findings from inside-out patch-clamp recordings indicate that zinc directly inhibits mSlo3 channels in a dose-dependent manner (Fig. 2). Zinc inhibition in mSlo3 channels is dependent on pH (Fig. 2A-E), voltage (Fig. 2G-H; Fig.2—figure supplement 1B, C) and exhibits a long-lasting inhibitory effect (Fig. 2I, K). This sustained inhibitory effect of zinc can be partially restored by EGTA, a divalent ion chelator with high affinity for zinc (Fig. 2J, K; Fig.2—figure supplement 1D, E). Previous studies indicate that zinc also regulates the Slo channel family, albeit with varying effects among Slo3 channel subtypes: zinc activates Slo1 channels (Hou et al., 2010) while inhibiting Slo2.2 channels (Budelli et al., 2016; J. Zhang et al., 2023). Zinc may produce these opposite effects on Slo2.2 and Slo1 by binding to distinct, non-conserved sites among the subtypes (J. Zhang et al., 2023).

Zinc inhibition on mSlo3 channels is characterized by the slow dissociation of zinc from the channel (Fig. 2I-K). Similar slow dissociation has been observed in zinc-regulated ion channels and receptors, such as NMDA and glycine receptors (Paoletti et al., 1997; Trombley et al., 2011). In NMDA receptors, high zinc affinity results in slow dissociation, which has been suggested to have a physiological relevance in neurons (Paoletti et al., 1997).

Slow dissociation typically indicates high-affinity binding. In contrast to NMDA receptors, which exhibit IC_50_ values ranging from nM to μM, mSlo3 has a relatively high IC_50_ for zinc inhibition (145.2 µM). This might be due to the ∼40-second zinc application time in our study, which may have been insufficient to fully observe the effects at lower concentrations (Fig. 2B, C). Extended exposure of intracellular zinc in sperm could potentially result in greater inhibition.

Furthermore, our results also show that when mSlo3 is coexpressed with Lrrc52 the lower concentration of zinc was sufficient to inhibit mSlo3 channels (Fig. 3). Although we still did not identify the exact mechanism underlying it, Lrrc52 shifts the gating of mSlo3 to more hyperpolarized potentials and lower pH, making the channel easier to open (Leonetti et al., 2012; Yang et al., 2011; X. H. Zeng et al., 2015). Considering that the zinc inhibition of Slo3 depends on membrane potential and intracellular pH, it is understandable that the presence of this auxiliary subunit would lead to more efficient inhibition by zinc.

### Structural determinants of zinc inhibition in the mSlo3 channel

We identified potential key amino acids in the mSlo3 channel that may serve as zinc binding sites using MD simulations and amino acid mutagenesis (Fig. 4, 5; Fig. 4—figure supplement 1, Fig. 5—figure supplement 1). Previous studies on the Slo channel family indicates that zinc binding sites are located in RCK1 domain for Slo1 (Hou et al., 2010) and RCK2 domain for Slo2.2 (J. Zhang et al., 2023). However, in mSlo3 channel, our results didn’t show that either RCK1 or RCK2 may serve as zinc binding site (Fig. 4— figure supplement 1).

On the other hand, our analysis of current recovery for mSlo3 mutants in the VSD region revealed that E169A and E205A showed significant recovery upon wash-out with pH 8.0. This result aligns with MD simulations showing faster zinc dissociation for both E169A and E205A, indicating their contributions in zinc-mediated inhibition. Thus, the cluster of negative residues within the VSD serve as the zinc binding site in the mSlo3 channel.

Based on sequence alignment, E169 (mSlo3 numbering) is conserved among Slo family channels in mice, whereas E205 (mSlo3 numbering) is not (Fig. 5—figure supplement 1F). To date, no studies have examined the corresponding residues to E169 (E191 in mSlo1 or E176 in mSlo2.2) for their potential zinc sensitivity, likely because the established zinc binding sites in these channels are located in the RCK1 domain for Slo1 (Hou et al., 2010) and the RCK2 domain for Slo2.2 (J. Zhang et al., 2023). The identified zinc binding site in Slo2.2 is conserved in Slo2.1 but is absent in both Slo1 and Slo3 (J. Zhang et al., 2023), further suggesting that zinc regulation differs among Slo family members. Although regions surrounding E191 or E176 may still provide additional insights into zinc regulation and could be of interest for future investigation, E205 stands out because, unlike E169, it is not conserved across the Slo family, making it unique to mSlo3 and potentially a specific determinant of zinc sensitivity in this channel.

However, our single amino acid mutations in this site, which contain clustered negative residues, did not significantly change zinc-mediated current reduction compared to the wildtype. This may be due to mutating one single amino acid might not be sufficient to fully elucidate other contributing residues in the predicted mSlo3 zinc binding site. Thus, more extensive mutagenesis studies will be needed to fully understand zinc inhibition in mSlo3.

It is worth noting that the incomplete loss of zinc sensitivity in these mutants suggests that additional mechanisms may participate in zinc modulation of Slo3. These may include modulation of nearby charged residues, structural rearrangements influenced by zinc binding, or the presence of multiple zinc binding sites. Comparisons with Slo2.2 (J. Zhang et al., 2023), KCNQ4 (Gao et al., 2017), and voltage-gated calcium channels (Sun et al., 2007) further support the possibility of diverse molecular determinants for zinc inhibition. Our VCF, mutagenesis, and simulation data together indicate that zinc influences voltage sensor movement in mSlo3, which may suggest a distinct inhibitory mechanism that warrants further investigation.

### Physiological relevance of zinc inhibition of the mSlo3 channel in mouse sperm

Zinc is thought to be involved in capacitation and the acrosome reaction, suggesting its role in regulating sperm fertilizing ability *in vitro* (Aonuma et al., 1978). This finding is supported by studies indicating that seminal zinc inhibits sperm’s ability to undergo the acrosome reaction (Riffo et al., 1992). However, the specific function of intracellular zinc in relation to the capacitation process has not been thoroughly investigated. Previous research indicates that zinc inhibits sperm motility in ejaculation medium as well as acrosome reaction (Henkel et al., 1999; Riffo et al., 1992), leading to the hypothesis that zinc might influence capacitation.

Slo3 plays a critical role in sperm fertility, since mice lacking Slo3 are infertile despite their normal sperm morphology (Santi et al., 2010). Furthermore, our previous work showed that VSP-deficient sperm exhibited enhanced Slo3 channel activity, which indirectly increases Ca^2+^ influx and results in abnormal sperm motility (Kawai et al., 2019). Taken together, our findings suggest that intracellular zinc is potentially a key regulator of sperm capacitation, functioning through inhibition of the Slo3 channel. Although these results support a mechanistic link between zinc and Slo3 activity, future studies that combine genetic or pharmacological modulation of Slo3 with comprehensive capacitation analyses will be required to define its physiological impact in more detail. Within this context, this study highlights the potential importance of intracellular zinc in the regulation of sperm capacitation.

In addition to that, to date, there are only few reports on the effect of zinc on other sperm ion channels, and none have been reported in mouse sperm. One important study was reported by (Jeschke et al., 2021), in which seminal zinc was found to inhibit prostaglandin-induced activation of CatSper, a sperm-specific Ca^2+^ channel, in human sperm. The complex opposing action of seminal zinc and prostaglandins on CatSper may help preventing premature activation of CatSper in the ejaculate and act as a dilution sensor, although this study does not provide direct evidence for zinc acting directly on CatSper (Jeschke et al., 2021).

Taken together, this study provides new insights into the zinc-dependent modulation of sperm ion channels, underscoring the central role of Slo3 and offering a foundation for future exploration of the complex molecular mechanisms governing fertilization.

## Materials and Methods

### Ethical approval

All animal experiments were approved by the Animal Care and Use Committee of the Osaka University Graduate School of Medicine and performed in accordance with its guidelines.

### Drug and Reagent

Zinc pyrithione was purchased from Fujifilm Wako (Osaka, Japan). TPEN was obtained from Cayman (Ann Arbor, MI, USA). Gramicidin was obtained from Merck (Kenilworth, NJ).

### Patch Clamp Recordings from Spermatozoa

Patch clamp recordings from spermatozoa were conducted as previously described (Kawai et al., 2019; Kirichok et al., 2006). Briefly, sperm were harvested from the mouse corpus epididymis using an HS-based solution containing (in mM): 135 NaCl, 5 KCl, 2 CaCl_2_, 1 MgSO_4_, 20 HEPES, 5 glucose, 10 lactic acid, and 1 sodium pyruvate (pH 7.4). After 10 minutes, the supernatant was collected, centrifuged, washed twice with the HS-based solution, resuspended, and placed on untreated glass coverslips.

For conventional whole-cell recordings, the intracellular solution contained (in mM): 120 KCl, 3 MgCl_2_, 40 MES, and 0.3 EGTA (pH 6.0), with the osmolality adjusted to approximately 300 mOsm/kg using sucrose. The same HS-based solution was used as the bath solution.

For perforated patch recordings, the intracellular solution included (in mM): 120 KCl, 3 MgCl_2_, 40 HEPES, 0.3 EGTA (pH 7.0), and 0.05 mg/ml gramicidin. HS-based solution was used as the extracellular solution. Once pores formed, the access resistance ranged between 50 and 100 MΩ.

Recording pipettes were made with borosilicate glass (BF-150-86-10; Sutter Instruments, CA, USA) using a puller (P-97; Sutter Instruments). The recordings were conducted using an Axopatch 200B amplifier (Molecular Devices, CA, USA) and sampled at 5 kHz using a Digidata 1550A system (Molecular Devices) with pCLAMP 10.5 software (Molecular Devices). Low pass filter of 20 Hz was applied to increase S/N ratio.

### Zinc imaging from Spermatozoa

Sperm were isolated from the cauda epididymis of mice and transferred to HS medium. The sperm were then loaded with 10 μM FluoZin3-AM (Invitrogen) and 0.05% Pluronic F-127 in HS medium, followed by a 30-minutes incubation at 37ºC. After incubation, the sperm were washed with HS medium and incubated on Cell-Tak (Corning, Bedford, MA)-coated round cover glasses for 10 minutes. The cover glass was mounted on a perfusion chamber for imaging experiments.

Fluorescence signals were detected using a filter set (Ex 495/10; DM 515; Em 535/25) with an IX-81 upright microscope (Olympus, Tokyo, Japan) equipped with a C9100 EMCCD camera (Hamamatsu Photonics, Shizuoka, Japan) and Metamorph imaging software (Molecular Devices). Images were captured every 30 seconds. Fluorescence intensity is measured by the signal taken from the whole head and the proximal part of tail in sperm.

For capacitation, TYH medium (Toyoda et al., 1971) was used, consisting of: 120 mM NaCl, 4.8 mM KCl, 1.2 mM KH_2_PO_4_, 5.6 mM glucose, 1.0 mM sodium pyruvate, 1.7 mM CaCl_2_, 1.2 mM MgSO_4_, 25 mM NaHCO_3_, 4.0 g/L ALBMAX I (Thermo Fisher Scientific), and penicillin (50 units/ml)-streptomycin (50 μg/ml).

The experiment focused on the head and tail regions of spermatozoa. For analysis, cells in which both the head and tail were visible throughout the entire experiment were selected.

### Sperm motility analysis

Sperm velocity was analyzed as previously described (Miyata et al., 2021). Briefly, spermatozoa isolated from the cauda epididymis were suspended in TYH medium. The TYH medium contained: 120 mM NaCl, 4.8 mM KCl, 1.2 mM KH_2_PO_4_, 5.6 mM glucose, 1.0 mM sodium pyruvate, 1.7 mM CaCl_2_, 1.2 mM MgSO_4_, 25 mM NaHCO_3_, 4.0 g/L ALBMAX I (Thermo Fisher Scientific), penicillin (50 units/ml) - streptomycin (50 μg/ml), and 0.6% Phenol Red.

Average path velocity (VAP), curvilinear velocity (VCL), straight-line velocity (VSL), straightness (STR), and linearity (LIN) was measured using the CEROS II sperm analysis system (Hamilton Thorne Biosciences, MA, USA) at 10 minutes and 2 hours after incubation. In some experiments, 20 μM TPEN was directly added to the TYH medium.

### Molecular Biology

All Slo3 mutants were constructed from WT *mus musculus* Slo3 (mSlo3) in PSD64TF. Single amino acid mutations were introduced using PrimeSTAR Mutagenesis Basal Kit (Takara Bio Inc.). Chimeras were generated using In-Fusion HD Cloning Kit (Takara Bio Inc.). The introduced mutations and generated chimeras were confirmed by DNA sequencing. The mMESSAGE SP6 RNA transcription kit (Thermo Fisher Scientific) was used to transcribe WT, mutants, and chimeras of mSlo3 cRNAs from plasmid cDNA linearized by Xba1 restriction enzyme (Takara Bio Inc.).

### Preparation of *Xenopus* laevis oocytes

As an anesthetic agent, 0.15% tricaine (Sigma-Aldrich / Tokyo Chemical Industry) was used for *Xenopus* laevis before surgical operation for isolation of oocytes. Follicular membranes were removed from isolated oocytes by collagenase treatment (2 mg ml−1; type 1; Sigma-Aldrich or Collagenase-P from Roche) for 3 hr. Oocytes were then rinsed and stored in ND96 solution (96 mM NaCl, 2 mM KCl, 1.8 mM CaCl_2_, 1 mM MgCl_2_, and 5 mM 2-[4-(2-Hydroxyethyl)-1-piperazinyl]ethanesulfonic acid (HEPES), pH 7.5, with NaOH) supplemented with 0.1 mg/mL gentamycin (FUJIFILM Wako Pure Chemical Corporation or Nacalai Tesque, Inc.) and 5 mM sodium pyruvate. The incubation temperature was 18 °C.

### Expression of mSlo3 channel

*Xenopus* oocytes injected with 15 ng of WT mSlo3 cRNA and incubated for 2 days at 18°C. For mSlo3 mutants, oocytes were injected with 50 ng–60 ng of cRNA and incubated for 1–3 days, depending on the desired expression level for patch-clamp and voltage-clamp fluorometry (VCF) experiments. cRNA for mLrrc52 was also injected for data shown in Fig. 3, Fig. 3—figure supplement 1, Fig. 6, and Fig. 6—figure supplement 1F-H.

### Oocyte patch-clamp recording of mSlo3

Patch-clamp recordings of *Xenopus* oocytes were performed using an Axopatch-200B amplifier (Molecular Devices) connected through an AD/DA converter (Digidata 1440A: Molecular Devices) and operated with pClamp software. Current traces were filtered at 5 kHz and sampled at 100 kHz. Vitelline membrane was manually removed using forceps after incubation for 1 to 5-minute in a hypertonic solution containing 192 mM NMDG, 4 mM KCl, 3.6 mM CaCl_2_, 2 mM MgCl_2_, and 10 mM HEPES, pH 7.4, with HCl. Leak currents were subtracted by using a P/-8 protocol for step pulses recordings only. Patch pipettes, with resistance ranging from 1 to 3 MΩ were pulled from borosilicate glass (Drummond Scientific Company). The pipette solution contains 0.5 mM potassium gluconate, 0.5 mM KCl, 1.1 mM KOH, 10 mM HEPES, 159 mM sodium gluconate, and 2 mM MgCl_2_, pH 7.1, adjusted with methanesulfonic acid.

Patches were perfused with: (1) bath solution containing 184 mM potassium gluconate, 10 mM KOH and 10 mM HEPES in pH 6.0, 7.5, and 8.0; (2) zinc-containing bath solution at pH 7.5 and pH 8.0, with zinc concentrations of 0.1, 1, 10, 100, and 1000 µM. (3) bath solution containing 5 mM EGTA in pH 8.0. All pH values were adjusted using methanesulfonic acid. Slo3 macroscopic currents were evoked by an alkaline pH and by a depolarizing ramp pulse (from -100 mV to +100 mV with the holding potential of either - 60 mV or 0 mV) or step pulses (from -100 mV to +180 mV with the holding potential at - 60 mV). All bath solutions were perfused using a high-speed perfusion system ALA-VM8 OCTAFLOW (ALA Scientific Instruments Inc.)

### Voltage-clamp fluorometry (VCF) recording of mSlo3

In mSlo3 VCF construct, the target labeling site (in S3-S4 linker) was substituted with cysteine residue. Two native, extracellular cysteines were mutated to avoid non-specific labeling (mSLo3 C19S/C131S). Oocytes for VCF recordings were labeled with Tetramethylrhodamine-5-maleimide (TMRM) (Invitrogen). In prior to VCF recordings, oocytes were incubated in a labeling solutions (120 mM K-methanesulfonate (MES), 2 mM CaCl_2_, and 10 mM HEPES pH 7.4) containing 10 µM TMRM (Savalli et al., 2006) on ice for 1 hour. Labeled oocytes were then rinsed in TMRM-free ND96 solution four times and kept in the dark until and throughout the recording.

Oocytes were placed in a recording chamber with the animal pole facing upward and held at -60 mV. For screening experiments, step pulses ranging from -80 mV to +160 mV were applied, while pulses from -160 mV to +160 mV were used for direct zinc injection experiments. For the VCF recording in the absence and presence of zinc, voltage steps were first applied to record the initial response. Subsequently, a puff of 10 mM zinc solution was manually injected into the same oocyte using a glass needle and positive pressure. Sixty seconds post-injection, a second set of step pulses was applied to record the response under zinc application. In VCF recording, borosilicate glass capillaries (Harvard Apparatus) were used with a resistance of 0.2–0.4 MΩ when filled with 3 M KOAc and 10 mM KCl. Bath solution contained 96 mM NaCl, 2 mM KCl, 1.8 mM CaCl_2_, 1 mM MgCl_2_, and 5 mM HEPES, pH 7.4-7.5, with NaOH was used for recordings.

Fluorometric recordings were performed with an upright fluorescence microscope (Olympus BX50WI) equipped with a water immersion objective lens (Olympus XLUMPLAN FL 20x/1.00) to collect the emission light from the voltage-clamped oocytes.

The light from LED source (MCWHL8 Thorlabs) was applied through a band-pass excitation filter (531 nm/40 nm BrightLine: Semrock). Emitted light was passed through band pass emission filter of 593nm/40nm (BrightLine, Semrock). The emission signals were detected by a photomultiplier (H10722-20; Hamamatsu Photonics). The detected emission intensities were acquired by a Digidata 1440A (Axon Instruments) and Clampex 10.3 software (Molecular Devices) at 25 kHz for VCF mutants screening and 50 kHz for VCF recording in the absence and presence of zinc. To improve the signal-to-noise ratio, VCF recording was repeated 3 times. Averaged data were used for data presentation and analysis.

### Three-dimensional structural modeling of mouse Slo3 and prediction of zinc binding site

The amino acid sequence of mouse Slo3 (mSlo3) (O54982-1) was obtained from the UniProt database (Fig. 4) and was retrieved from NCBI NP_032458.3 (Fig. 5). The sequence (only the transmembrane region from residue M1 to G330, or the full-length protein) was input to the AlphaFold3 (Abramson et al., 2024) structural modelling web platform to generate an initial structure with (Fig. 4A) or without (Fig. 5A) 50 zinc ions. All the structural data presented in this study were generated using PyMOL molecular graphics system ver. 2.5.0 (Schrodinger LLC).

### Flooding simulations

The full-length mSlo3 from the AlphaFold3 webserver was trimmed to remove long disordered regions (residue 1-18, 616-698 and 1050-1121), resulting in 12 separate chains. The mSlo3 complex was embedded into the POPC bilayer and solvated in 0.15 M NaCl and 0.15 ZnCl_2_ using CHARMM-GUI Membrane Builder (Wu et al., 2014). The production run simulations were simulated with a 2 fs timestep using the CHARMM36m forcefield (Huang et al., 2016) in GROMACS 2023.3 (Abraham et al., 2023). The system was then energy minimized and equilibrated using the standard six steps CHARMM-GUI equilibration protocol. This includes the following set-up: The protein backbone was restrained at the force constant of 4000, 2000, 1000, 500, 200 and 50 kJmol^-1^nm^-2^, the protein side chains were restrained with the force constant of 2000, 1000, 500, 200, 50, and 0 kJmol^-1^nm^-2^, the lipids non-H atoms were restrained at the force constant of 1000, 400, 400, 200, 40 and 0 kJmol^-1^nm^-2^ and the dihedral restraint was set at the force constant of 1000, 400, 200, 200, 100 and 0 kJmol^-1^rad^-2^. The simulations were equilibrated with a 1 fs timestep for 125 ps for the first three steps, and then to a 2 fs timestep for 500 ps the next two, and 5 ns for the final step. The first two steps were conducted with the NVT ensemble, whereas the last four were conducted with the NPT ensemble. All equilibration runs were conducted at 310 K using Berendsen thermostat (Berendsen et al., 1984). In all NPT ensemble equilibration, the semi-isotropic pressure was maintained at 1 bar using Berendsen barostat. The production runs were conducted for 200 ns under 310 K using the v-rescale thermostat (Bussi et al., 2007). The semi-isotropic pressure of all systems was maintained at 1 bar using a C-rescale barostat (Bernetti & Bussi, 2020). All simulations were carried out in triplicates, where velocity was randomly generated at the beginning of every simulation. All simulation frames were saved every 0.1 ns.

### Zn stability

Based on the flooding simulations, the binding pose for Zn^2+^ is adjusted so that the ion is 3-4 Å away from E169. This results in the system with 4 Zn^2+^ per system, one per subunit. The system was then energy minimized using the steepest descent algorithm and equilibrated with the Cα restraint at 1000 kJ kJmol^-1^nm^-2^ for 10 ns. The production run was conducted for 200 ns, with the same barostat and thermostat setting as the flooding simulation.

### Data Analysis

Inside-out patch-clamp data were analyzed using Clampfit 11.3 software (Molecular Devices), Igor Pro 6.22A (Wavemetrics), and GraphPad Prism 10. Analysis of current-voltage (I-V) were obtained from the maximum current amplitude at steady-state. Normalization for figure 2H was performed by dividing the absolute current of mSlo3 at pH 8.0 of each voltage by the absolute current at the pre-determined highest voltage that still produced a stable mSlo3 current (i.e., good patch, good clamp). In this analysis, +140 mV was chosen as the highest voltage for normalization, since in few cells the patch was lost at +160mV and +180mV. Similar to the control condition, the absolute current of mSlo3 in the presence of 100 µM zinc was normalized to the absolute current of the control at +140 mV. I-V relationship was fitted with Boltzmann equation:

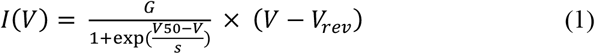

where G is a scaling factor (maximum conductance or current), V_50_ is the midpoint, and s is the slope value.

And then the I-V relationship in the presence of 100 µM of ZnCl_2_ was fitted with Woodhull equation (Woodhull, 1973) to express the current as a function of voltage, incorporating both the voltage-dependent gating of the channel and the voltage-dependent blocking of the channel:

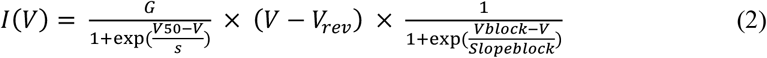

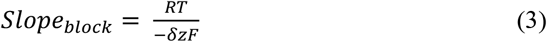

where V_block_ is the voltage at which blocking is half-maximal, Slope_block_ is the slope factor for the voltage dependence of the blocking effect, and δ is the fraction of the membrane potential that the blocking ion experiences.

V_rev_ is the reversal potential and was calculated with following equation as described in (Schreiber et al., 1998) using Goldman-Hodgkin-Katz equation:

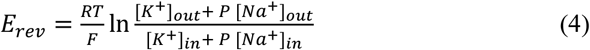

where P is the Na^+^ to K^+^ permeability ratio (Schreiber et al., 1998).

Conductance-voltage (G-V) relationship of Slo3 were obtained from conductance calculation using the following equation:

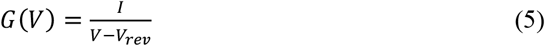

V_rev_ was calculated using equation (4) and fitted to Boltzmann equation in (1), adjusted for G-V.

IC_50_ (half maximal inhibitory concentration) was calculated using equation while constraining the Hill Slope to a constant value of -1.0:

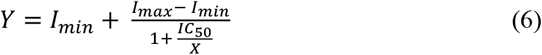

where I_max_ is the maximal current when inhibition is minimal, I_min_ is the minimal current inhibition is maximal, and X is the inhibitor concentration.

Arithmetic operations were performed by Clampfit to compensate bleaching from the fluorescence traces:

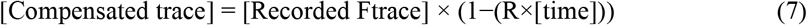

where [time] is the value of the point given by Clampfit.

All the compensated traces were then normalized by setting each baseline level (F signal at −60 mV, the holding potential) to be 1, to calculate the % F change:

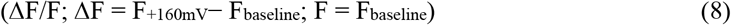

Conductance-voltage (G-V) relationship for VCF data were obtained from conductance calculation using equation (5). Fluorescence-voltage (F-V) relationship were obtained from the normalized ΔF/F and fitted with the equation (1).

### Statistical analysis

Statistical analysis was performed by either one-way ANOVA, unpaired t-test or paired t-test. Following one-way ANOVA, Dunnett’s post-hoc test was applied. The data were expressed as mean± s.e.m. with n indicating the number of samples. Values of p were stated in each bar graph. All the statistical analysis and the bar graphs were performed and generated with GraphPad Prism 10.

## Acknowledgments

The authors thank Prof. Fredrik Elinder (Linköping University, Sweden) for discussion and the guidance on analysis and experiments, Ms. Hikari Ginama and Ms. Megumi Kobayashi for technical support and Dr. Natsuki Mizutani for discussion and guidance of experiment. This work was supported by fundings provided by Grants-in-Aid from MEXT KAKENHI Grant Number JP20H05507 (T.K), JP20H05791 (Y.O), JP22K15074 (R.T.A), JSPS KAKENHI Grant Number JP17K15558, JP23K06334 (T.K), 19H03401, 22H02804 (Y.O), and JST FOREST Program Grant Number JPMJFR225Z (T.K). The research is supported by JSPS Postdoctoral Fellowship for Research in Japan (R.T.A) and Lee Kuan Yew Postdoctoral Fellowship from Nanyang Technological University, Singapore (T.P). Additionally, financial support was received from the Takeda Science Foundation and Mochida Memorial Foundation for Medical and Pharmaceutical Research, all of which contributed to T.K.’s research.

## Figures

**Figure S1—figure supplement 1.**
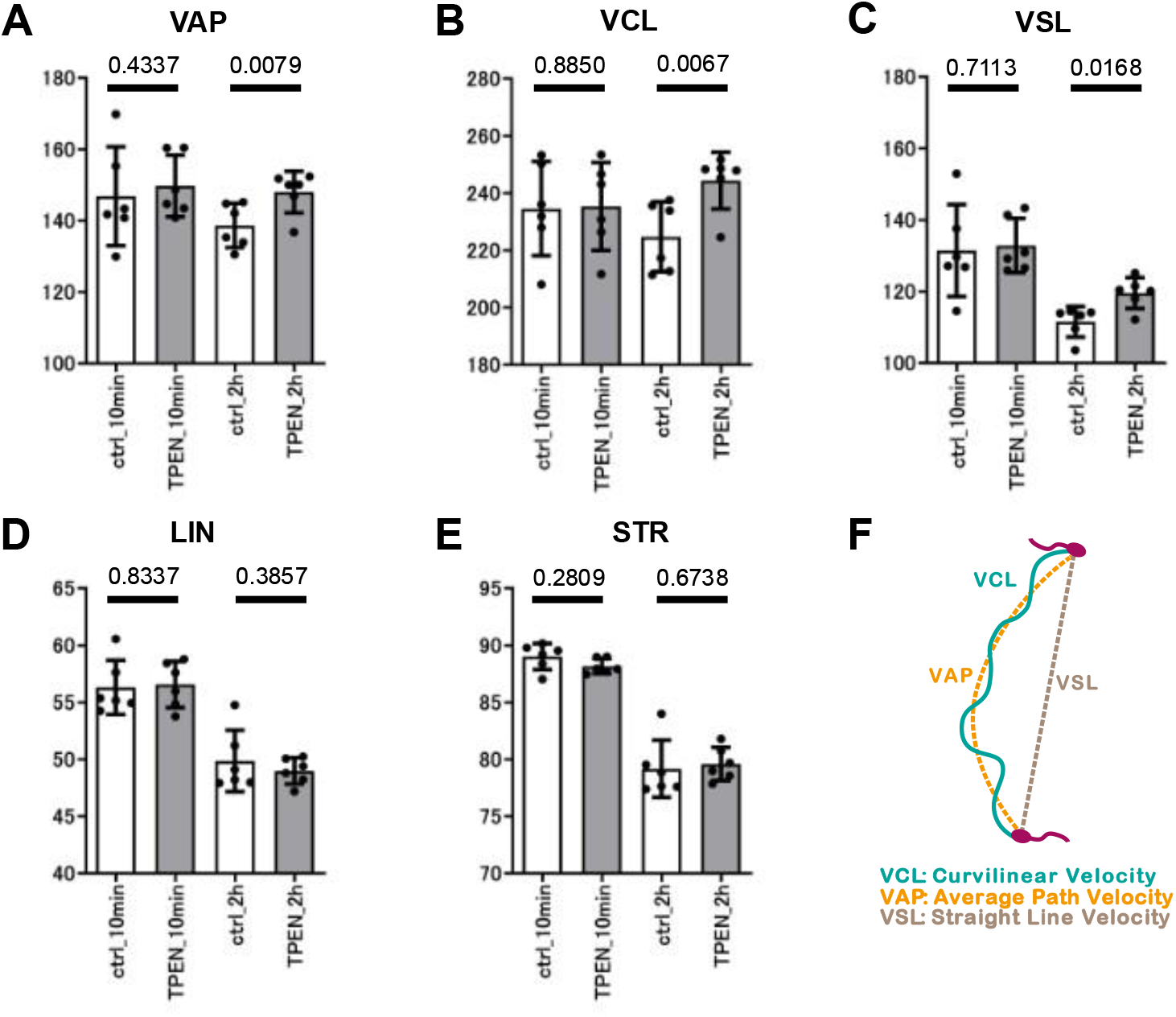
Effects of intracellular zinc on sperm motility before and after capacitation. Comparison of VAP, VCL and VSL show that intracellular zinc affects these kinds of motions only after the sperm underwent capacitation. (**A-E**) Quantitation of sperm motility parameters at: 10 min (control); 10 min in the presence of zinc chelator TPEN; 2 h after TYH incubation, a well-established capacitation-inducing medium (control); and 2 h after THY incubation in the presence of TPEN. Bar graph shows the comparison of (**A**) VAP (average path velocity); (**B**) VCL (curvilinear velocity); (**C**) VSL (straight-line velocity); (**D**) STR (straightness); and (**E**) LIN (linearity) as depicted in (**F**). All error bars are ± SD centered on the mean.

**Figure S1—figure supplement 2.**
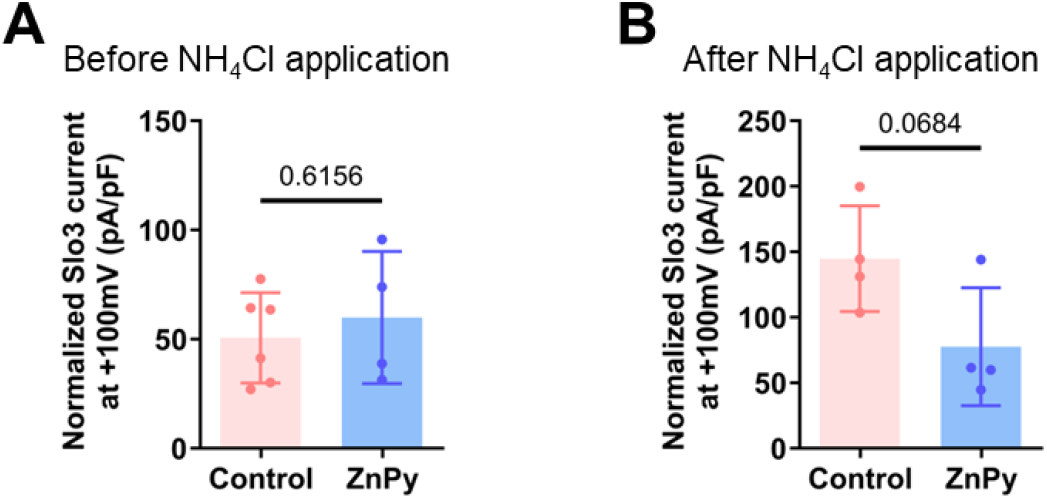
The effects of zinc on Slo3 current in sperm cells. (**A**) Comparison of normalized Slo3 current at +100 mV before NH_4_Cl application between control (50.67 ± 20.67 pA/pF; n=6) and in the presence of 100 µM ZnPy (59.95 ± 30.24pA/pF; n= 4), p=0.6156, Welch’s t-test. (**B**) Comparison of normalized Slo3 current at +100 mV after NH_4_Cl application between control (144.8 ± 40.36 pA/pF) and in the presence of 100 µM ZnPy (77.59 ± 44.97pA/pF; n= 4), p=0.0684, Welch’s t-test. All error bars are ± SD. centered on the mean.

**Figure S2—figure supplement 1.**
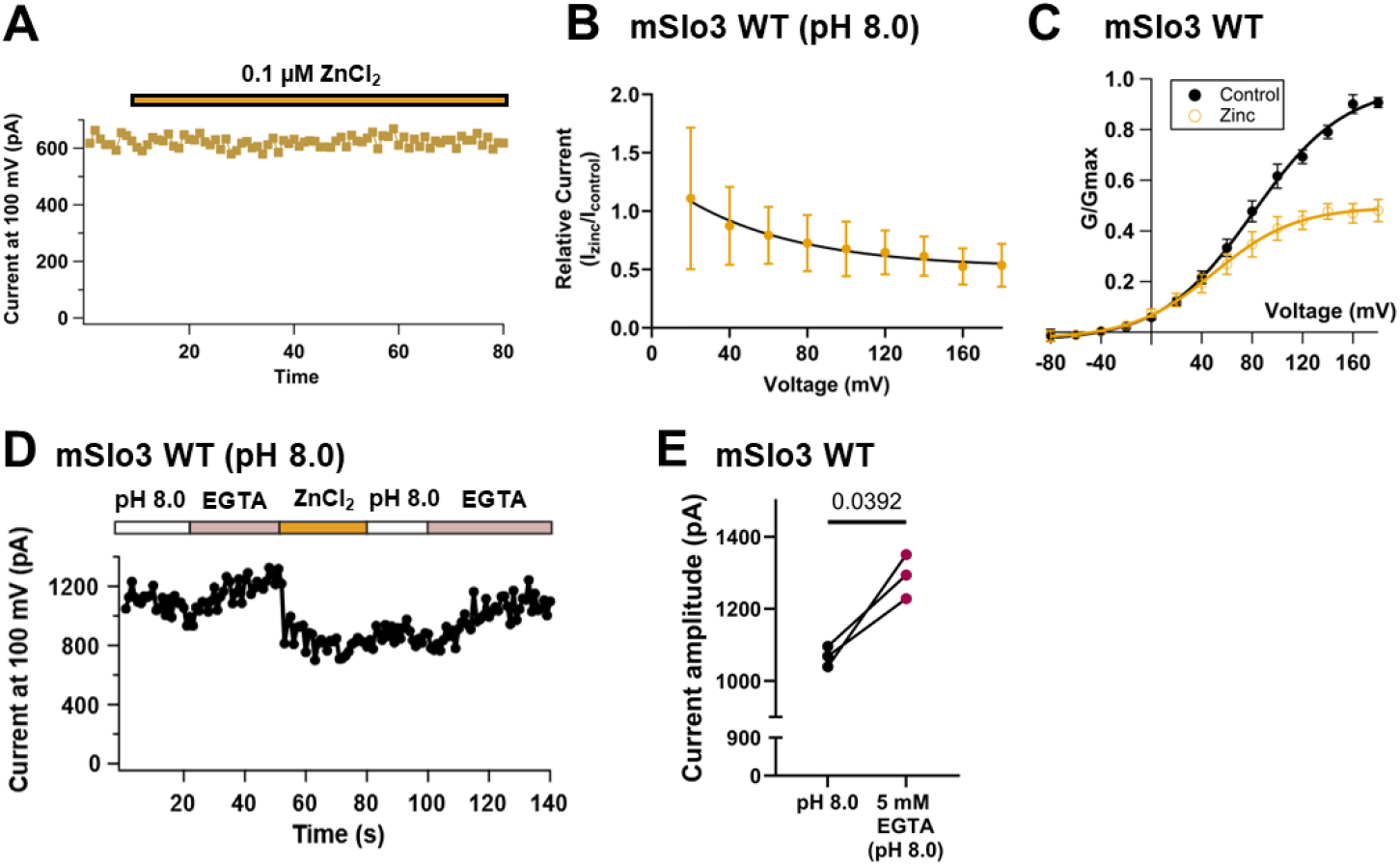
Zinc inhibits mSlo3 current. (**A**) Time course of Slo3 current at + 100 mV upon the application of 0.1 µM zinc. (**B**) Relationship between relative current (I_zinc_/I_control_) with membrane voltage at pH 8.0 shows that zinc inhibits mSlo3 current in voltage-dependent manner. (**C**) G-V relationship comparison of mSlo3 current between control (pH 8.0; black filled circle) and upon the application of 100 µM zinc in pH 8.0 (yellow orange circle) (V_50_ control = 80.67 ± 2.32; V_50_ zinc = 49.677 ± 1.38; n=13). Normalization was done based on the maximum conductance in control. (**D**) Time course of the change in current amplitude at +100 mV in response to pH 8.0, 5 mM EGTA in pH 8.0, 100 µM zinc, pH 8.0, and 5 mM EGTA in pH 8.0 as indicated (white: pH 8.0; yellow orange: 100 µM zinc in pH 8.0; pale lilac: 5 mM EGTA in pH 8.0). The baseline current showed a slight increase upon perfusing with 5 mM EGTA, indicating that endogenous zinc is already present in mSlo3, and it is affecting the baseline current. (**E**) Comparison of percentage of mSlo3 current evoked by pH 8.0 and 5 mM EGTA in pH 8.0 (p=0.0392). Statistical analysis was done by paired t-test, n=3. All error bars are ± SD centered on the mean.

**Figure S3—figure supplement 1.**
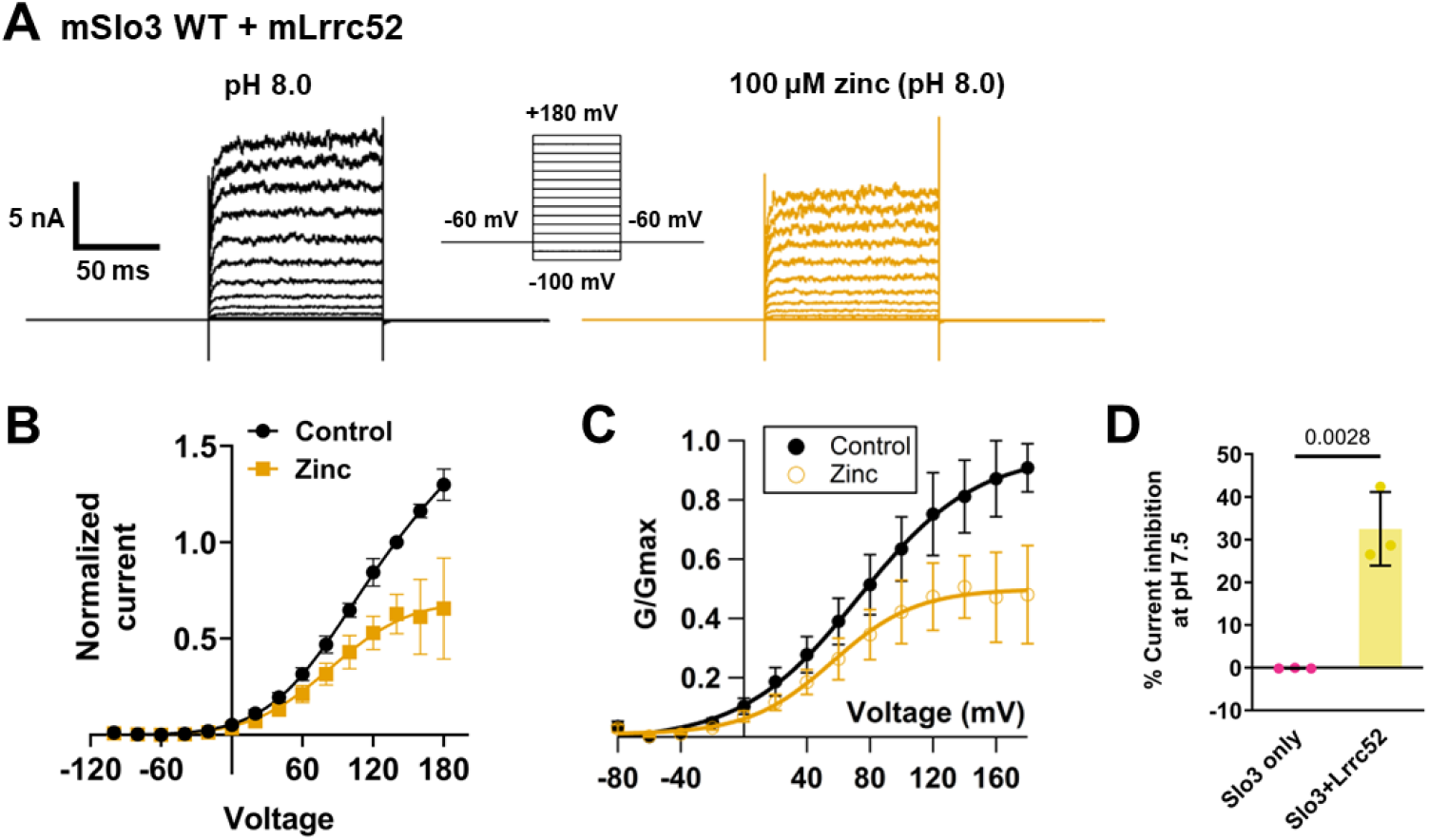
Zinc inhibits mSlo3 current when coexpressed with auxiliary Lrrc52. (**A-C**) Inside-out patch-clamp recording of mSlo3 co-expressed with mLrrc52 with pHi=8.0 by applying step pulses from –100 mV to +180 mV with the holding potential of -60 mV. Zinc inhibition is voltage-dependent. (**A**) Representative current traces upon the application of pH 8.0 (black) and 100 µM of zinc in pH 8.0 (yellow orange). (**B**) I-V relationship of control (pH 8.0; black) and zinc (100 µM; yellow orange). I-V curve was fitted by Boltzmann equation for control and followed by Woodhull equation for zinc inhibition (V_50 Ctrl_ = 74.01; Slope _Ctrl_ = 36.47; V_block_ Zn = 185.5; Slope_block_ Zn = -130.4; n=8). (**C**) G-V relationship comparison of mSlo3 current co-expressed with mLrrc52 between control (pH 8.0; black filled circle) and upon the application of 100 µM zinc in pH 8.0 (yellow orange circle) (n = 13). V_50_ control = 72.93 ± 2.06 V_50_ zinc = 56.22 ± 2.75. Normalization was done based on the maximum conductance in control. (**D**) Comparison of the percentage of Slo3 current inhibition at pH 7.5 between Slo3 only (-0.13 ± 0.08 %; n=3) and Slo3+Lrrc52 (32.51 ± 8.62 %; n=3), p=0.0028, unpaired t=test. All error bars are ± SD centered on the mean.

**Figure S4—figure supplement 1.**
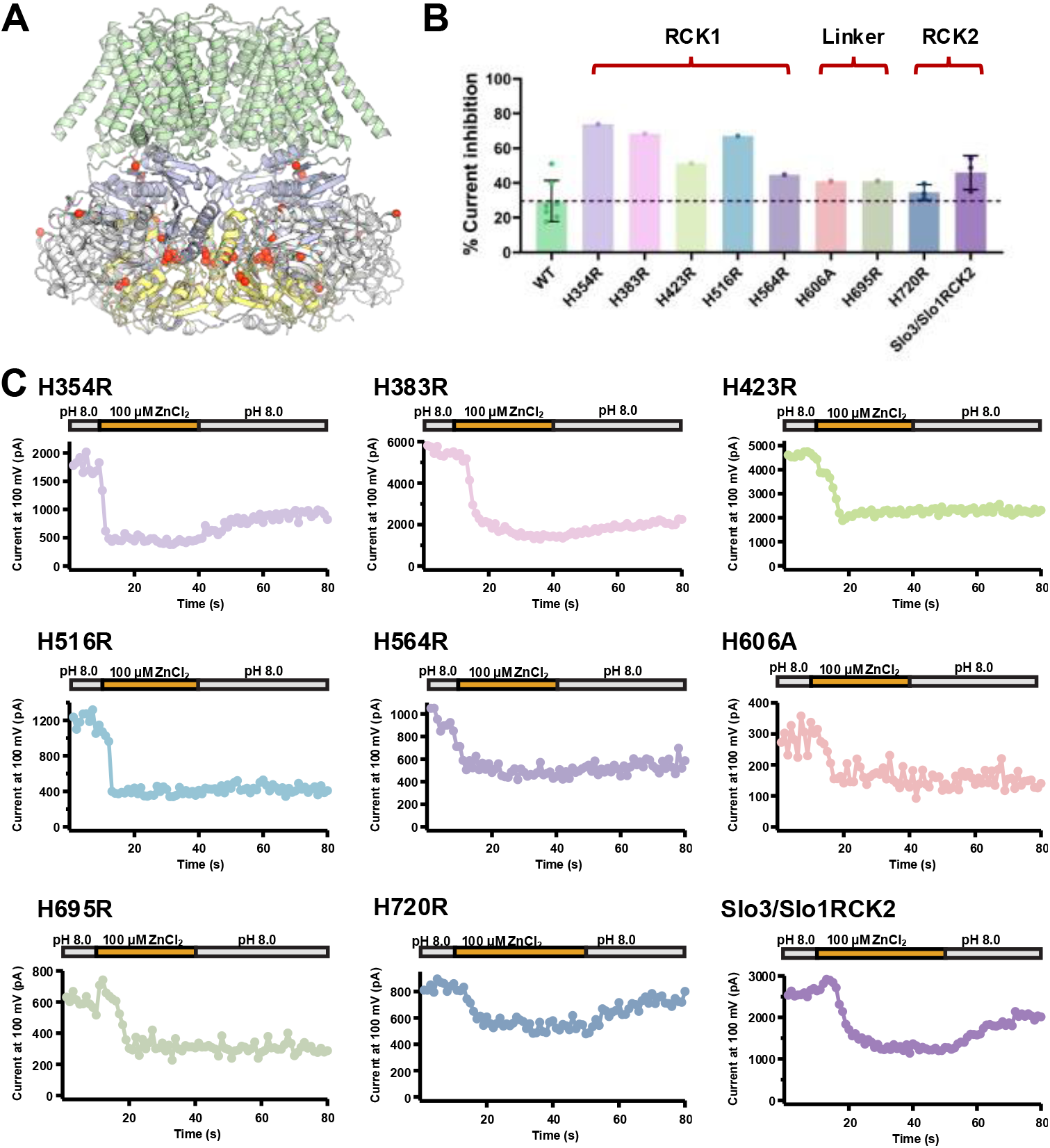
Computational studies and scanning mutagenesis of zinc binding site in mSlo3 channel. (**A**) The full-length Slo3 tetramer structure predicted by AlphaFold3, with RCK1 and RCK2 represented in blue and yellow cartoons, respectively, and 50 Zn^2+^ ions depicted as red spheres.(**B**) Comparison of the percent current inhibition upon the application of 100 µM zinc between wildtype and all the histidine mutants scanned (histidine residues located in RCK1, RCK1/RCK2 linker, RCK2, and the chimera in which the RCK2 of mSlo3 is replace with the RCK2 from mSlo1). Dotted black line indicates mean of control (WT). (**C**) Time course of the change in current amplitude at +100 mV in response to pH 8.0, 100 µM zinc, and pH 8.0 for wash-out as indicated for all the mutants as stated. All error bars are ± SD centered on the mean.

**Figure S5—figure supplement 1.**
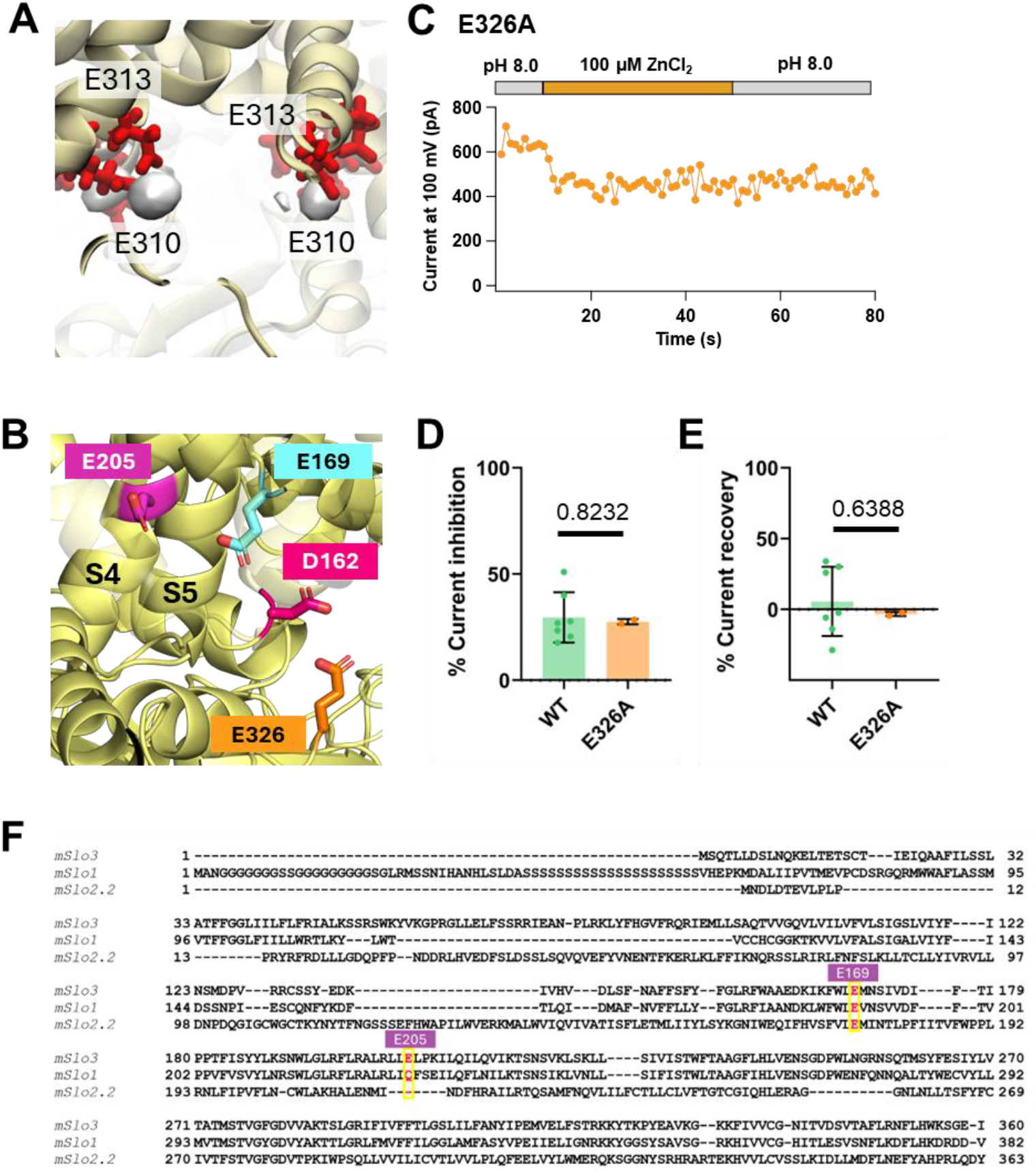
Computational studies and mutagenesis of zinc binding site in mSlo3 channel. (**A**) The density of Zn ions within 4 Å of the protein molecules was calculated using VolMap averaged across 200 ns flooding simulations (n = 3). Key acidic contacting residues are shown as red sticks. (**B**) Tetrameric mSlo3 structure representation of selected region which indicates D162 (S2-S3 linker), E169 (S3 domain), E205 (S4 domain), and E326 (intracellular linker between TM and RCK1) which have been mutated to elucidate the zinc binding site in mSlo3. (**C**) Time course of the change in current amplitude at +100 mV in response to pH 8.0, 100 µM zinc, and pH 8.0 for wash-out as indicated for E326A. (**D**) Comparison of the percent current inhibition upon the application of 100 µM zinc between wildtype and E326A (p=0.8232, unpaired t-test, n=2-7). (**E**) Comparison of percentage of current recovery upon wash-out using pH 8.0 between WT and E326A (p=0.6388, unpaired t-test, n=2-7). All error bars are ± SD centered on the mean. (**F**) Sequence alignment of mSlo3 (NP_032458.3), mSlo1 (NP_001240293.1) and mSlo2.2 (NP_001138875.1) using ClustalO. E169 and E205 residues (mslo3 numbering) are highlighted in red text with a yellow box.

**Figure S6—figure supplement 1.**
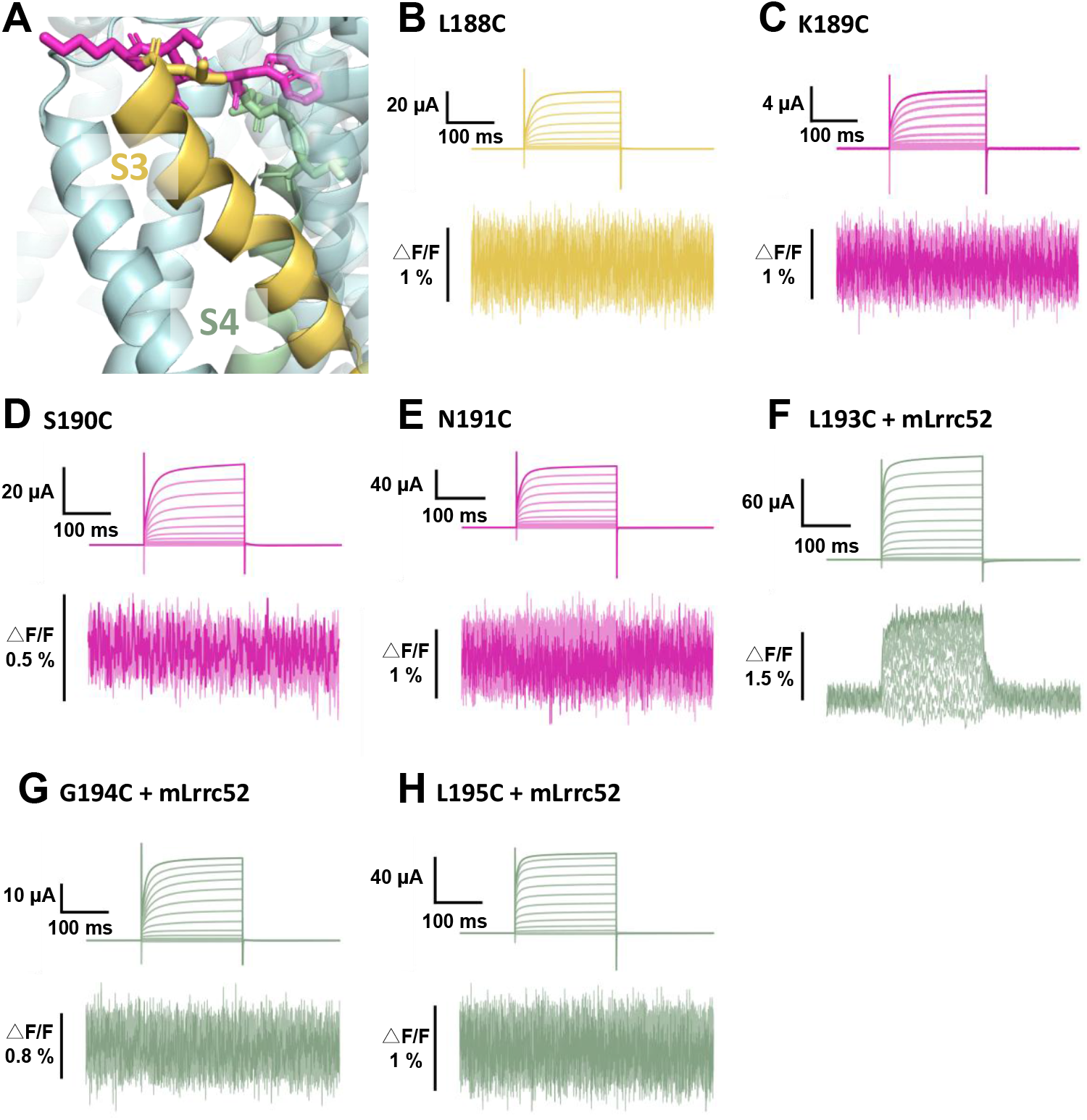
Scanning mutagenesis of mSlo3 channel VCF construct. (**A**) Structural representation of the VCF scanning regions in the mSlo3 channel, highlighting the top of S3 (yellow), the S3-S4 linker (violet), and the top of S4 (mud green). (**B-H**) Representative current traces and fluorescence signals recorded upon voltage steps from -80 mV to +160 mV with a holding potential of -60 mV for L188C, K189C, S190C, N191C, L193C (+mLrrc52), G194C (+mLrrc52), and L195C (+mLrrc52), respectively (n=4-7; ΔF/F mSlo3 L193C+Lrrc52: = 1.15 ± 0.48 % [n = 4]).

